# Information integration during bioelectric regulation of morphogenesis in the embryonic frog brain

**DOI:** 10.1101/2023.01.08.523164

**Authors:** Santosh Manicka, Vaibhav P. Pai, Michael Levin

## Abstract

Spatiotemporal bioelectric states regulate multiple aspects of embryogenesis. A key open question concerns how specific multicellular voltage potential patterns differentially activate distinct downstream genes required for organogenesis. To understand the information processing mechanisms underlying the relationship between spatial bioelectric patterns, genetics, and morphology, we focused on a specific spatiotemporal bioelectric pattern in the *Xenopus* ectoderm that regulates embryonic brain patterning. We used machine learning to design a minimal but scalable bioelectric-genetic dynamical network model of embryonic brain morphogenesis that qualitatively recapitulated previous experimental observations. A causal integration analysis of the model revealed a simple higher-order spatiotemporal information integration mechanism relating the spatial bioelectric and gene activity patterns, where the latter is expressed as a function of the causal influence of the voltages of groups of cells. Specific aspects of this mechanism include causal apportioning (certain cell positions are more important for collective decision making), informational asymmetry (depolarized cells are more influential than hyperpolarized cells), long distance influence (genes in a cell are variably sensitive to voltage of faraway cells), and division of labor (different genes are sensitive to different aspects of voltage pattern). The asymmetric information-processing character of the mechanism led the model to predict an unexpected degree of plasticity and robustness in the bioelectric prepattern that regulates normal embryonic brain development. Our *in vivo* experiments verified these predictions via molecular manipulations *in Xenopus* embryos. This work shows the power of using a minimal *in silico* approach to drastically reduce the parameter space *in vivo*, making hard biological questions tractable. These results provide insight into the collective decision-making process of cells in interpreting bioelectric pattens that guide large-scale morphogenesis, suggesting novel applications for biomedical interventions and new tools for synthetic bioengineering.

## Introduction

Embryonic development is a remarkable process, in which a large number of cells cooperate towards invariant large-scale anatomical shapes ^1,2^. Central to this phenomenon (in most organisms) is long-range order: not only local identity for each cell in a hardwired mosaic, but high levels of tolerance to noise, ability to handle perturbations, and active cell-cell interactions which maintain growth and form ^1,3,4^. Robust mechanisms ensure that overall proportions among organs, symmetry, and topological and geometrical relationships are maintained across the whole organism ^5–8^. It is especially important to understand context-sensitive responses that enable transcriptional events based on large-scale (non-cell-autonomous) information that signals completion of morphogenetic processes or errors in tissue-level order ^2,9^.

Morphogenesis is implemented by several physical modalities that enable cells to coordinate their actions toward specific target morphologies. In addition to biochemical gradients ^10–13^ and biomechanical forces ^14–16^, it is now evident that bioelectric networks ^17–19^ (spatiotemporal patterns of membrane voltage across cell fields) produce spatial gradient information that is instructive for growth and form. For example, endogenous bioelectric prepatterns have been functionally implicated in the location and size of organs such as the eye ^20^, invertebrate wing ^21–24^, vertebrate appendage ^25–27^, and face ^27–29^. Bioelectric prepatterns also polarize the left-right ^30–33^ and dorso-ventral ^34,35^ axes, and are involved in size control ^36–40^, embryonic compartmentalization ^41–45^, and stem cell function ^46–48^. Transduction mechanisms for bioelectric signals ^49^, downstream transcriptional machinery ^50^, and interfaces between electrical and mechanical events ^51,52^ have begun to be characterized. Moreover, computational models have now come online ^53–58^ that enable rational manipulation of ion channel and gap junction interfaces (via drugs, light, or mRNA misexpression) to induce ectopic organ formation ^20,59^, trigger complex regenerative response ^60–62^, and reverse tumorigenesis ^63,64^ in model systems *in vivo*. Some of these modulators are beginning to be applied to human medicine as electroceuticals ^65–74^.

While a number of mechanisms for converting voltage changes in an individual cell into changes of that cell’s behaviors have been characterized ^49^, a major gap in our knowledge concerns how *tissue-level* bioelectric information (distribution of cell resting potentials across cell fields) is processed for achieving large-scale morphological outcomes. For example, specific distributions of V_mem_ indicate the location, size, and organ identity of hearts, eyes, wings, and other structures ^20–22,75–77^. This requires downstream transcriptional cascades to be triggered by specific multicellular distributions of voltage. Because complex organogenesis cannot be solved by purely local information at the single cell level, it is essential to understand how the regional spatial pattern of voltages across a *group* of cells is converted into the diverse patterns of gene expression required for normal morphology ^1,9^. Thus, models of developmental bioelectricity must move to considering the more complex and interesting systems-level question of how a collective of cells processes patterns across distance to orchestrate a differentiated pattern of transcriptional responses ^55,57,78,79^. This in essence is the task of cracking the bioelectric code: mapping from voltage prepattern states to transcriptional patterns at a tissue-wide level ^80^.

The relationships between bioelectric gradients and downstream transcriptional cascades are not easily intuited, and the parameter space for experimentally testing all possibilities is vast. We therefore sought to establish and test a computational model of bioelectric pattern encoding in which we explicitly simulate bioelectrical states and gene-regulatory networks to understand their interplay. We chose to model the nascent *Xenopus laevis* brain, because it is a complex organ commonly used to model disorders with biomedical relevance ^81–87^, and because its normal morphogenesis has already been shown to include a bioelectric component ^20,88–90^. Briefly, beginning at embryonic stage 16, a dramatic contrast develops between the transmembrane voltage potential at the neural plate (hyperpolarized) and in the surrounding ectoderm (depolarized), establishing a bioelectric pre-pattern that regulates subsequent large-scale brain development ^88,89^ (Fig. 1A-B). Crucially, it is the long-range difference between voltage in these regions that is required for normal development, not the absolute values of specific regions. This bioelectric contrast pattern regulates expression of canonical brain development transcription factors (e.g., *Notch, Otx2, Pax6, Emx, Xbf1/Foxg1, Sox2*) as well as neural progenitor cell proliferation and apoptosis, driving normal development of the brain ^88,89,91^ (Fig. 1A-B).

**Figure 1.**
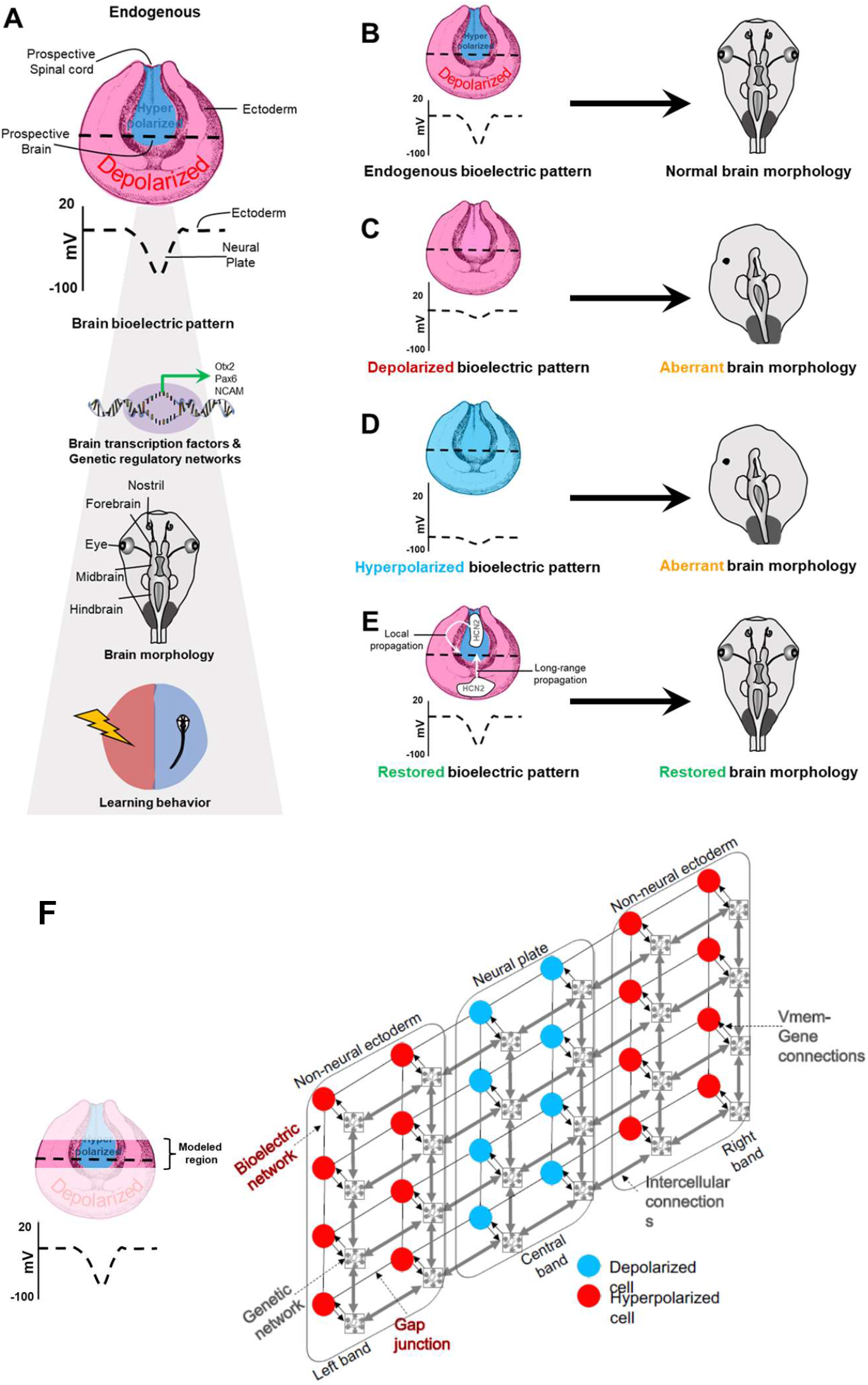
Schematic summary of endogenous voltage prepattern’s control of embryonic brain patterning in *Xenopus* and an overview of the neural plate circuit model of this process. (**A**) Summary of our previous studies ^88,89,91,93^ showing that the *difference* (dashed black line) in voltage across the hyperpolarized neural plate and relatively depolarized surrounding ectoderm is essential for driving correct gene expression pattern, large-scale brain morphology, and normal learning behavior. (**B**) Endogenous embryonic voltage prepattern resulting in normal brain morphology ^88,89,91,93^. (**C**) Embryonic manipulation of ion fluxes, exposure to teratogens (nicotine and ethanol), or genetic/biochemical disruption (Notch disruption) leading to depolarization of the neural plate erases the critical voltage difference between neural plate and ectoderm, resulting in mispatterned gene expression, and defects in large-scale brain morphology and learning behavior ^88,89,91–94^. (**D**) Embryonic manipulation of ion fluxes leading to hyperpolarization of surrounding ectoderm also erases the critical voltage difference between neural plate and ectoderm, resulting in mispatterned gene expression and abnormal large-scale brain morphology ^88,89^. (**E**) Both local (neural plate) or distant (ectodermal) interventions (manipulating ion fluxes by channel misexpression or drugs targeting ion channels) that restore this critical voltage difference between neural plate and ectoderm, restore gene expression, large-scale brain morphology, and learning behavior ^88,89,91–94^. (**F)** A schematic of our model. The overt structure of the model is a 2-dimensional lattice approximately representing the neural plate tissue (shown in the illustration). The circles indicate the individual cells of the tissue, with the colors indicating different voltages corresponding to the endogenous voltage pre-pattern (blue means depolarized and red, hyperpolarized). Each cell possesses a generic hyperpolarizing channel and depolarizing channel (not shown) and a generic gene regulatory network (grey networks enclosed in squares). The cells are connected both by gap junctions (thin black lines) that conduct the flow of ions, and by other intercellular connections (thick grey lines with arrowheads) conducive to the flow of gene products and proteins. The voltage and the gene network of a cell are bidirectionally connected to each other (thin black lines with arrowheads): the voltage acts as an external input to a subset of the genes, affecting their expression and in turn, the expressions of a subset of the genes can dynamically alter the conductance of the ion channels. The model is symmetrical in terms of the intercellular connectivity, that is, no directionality, such as that imparted by planar cell polarity, for example, is assumed. The 4×6 size of the lattice is accurate, as models of that size were used during training.

**Figure 2.**
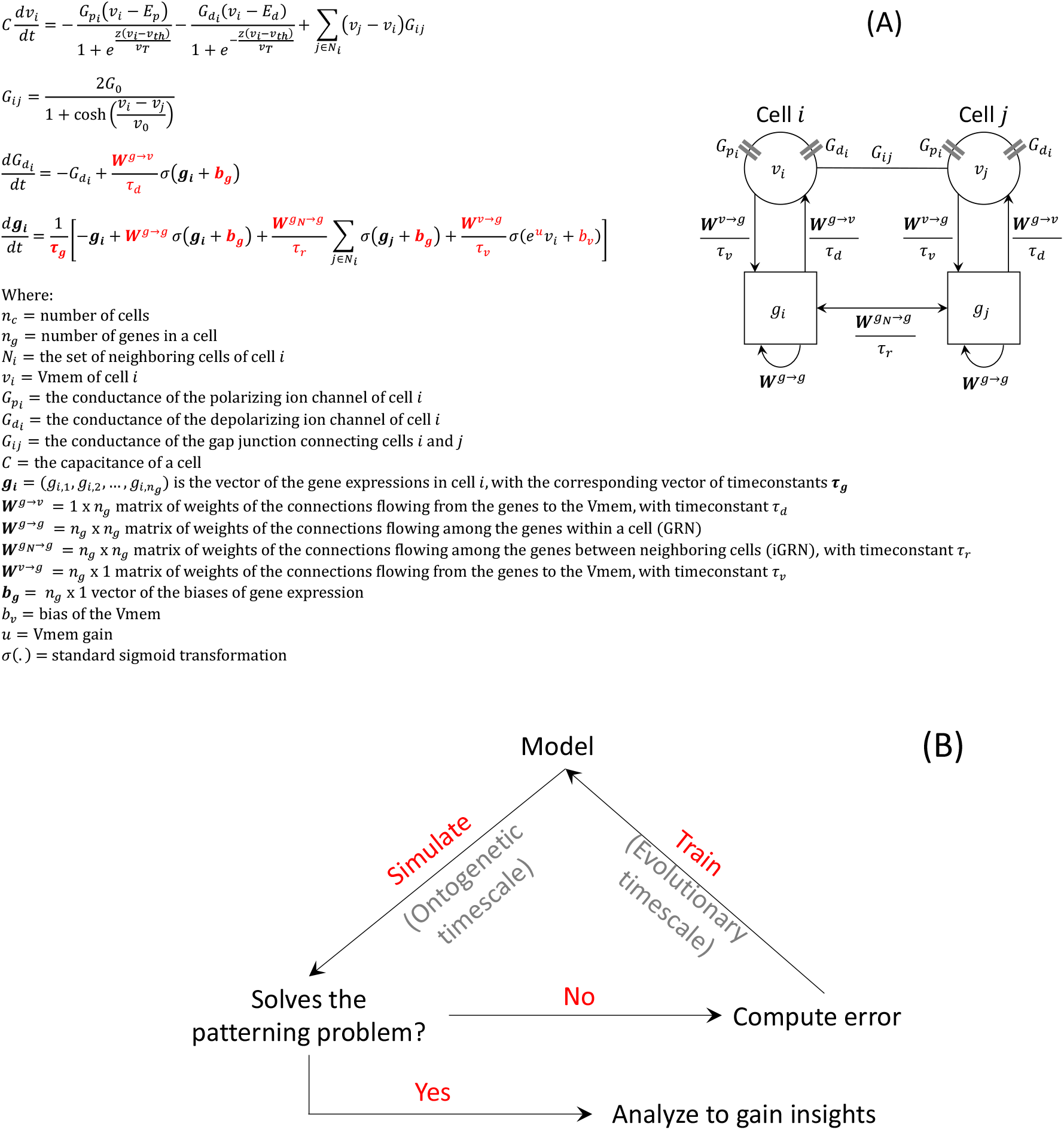
Overview of the training process of the neural plate circuit model. (**A)** The equations defining the model, with the variables and parameters mapped to a simplified two-cell version (the rest of the model is symmetrical to this version). The parameters (red) are learned using machine-learning techniques. The bioelectric constants, namely, *C, v_th_*, *v*_0_, *v_T_* and *G*_0_ are described in ref ^93^ and in the SI.1. **(B)** The model is trained using a combination of genetic algorithm and gradient descent. The final model referred to throughout this paper is a product of a training loop that starts with a population of randomly parametrized models, simulates all of them, and then modifies them using genetic algorithm and gradient descent to improve their ability to solve the pattern discrimination problem.

Disrupting this normal bioelectric contrast pattern disrupts the differential expression of these transcription factors, resulting in abnormal large-scale brain morphology and function ^88,89^ (Fig. 1C-D). In addition, neuroteratogens such as nicotine and ethanol and genetic or biochemical defects such as a disrupted notch signal cause severe brain morphology defects and impaired function by disrupting this bioelectric contrast pattern ^91–94^ (Fig. 1C-D). Remarkably, local and long-range interventions that restore the normal bioelectric pattern in the presence of these disrupters also rescue large-scale brain morphology and function, showing the incredible regulatory importance of this pattern in embryonic brain development ^88,91–94^ (Fig. 1E). Conversely, interfering with normal expression of Notch or Pax6 also disrupts the bioelectric contrast pattern, demonstrating feedback from gene expression to the large-scale bioelectric contrast pattern, at least for these two genes ^20,88,92^. Furthermore, recreating the bioelectric contrast pattern in other regions of the embryo is sufficient to re-specify those regions to produce ectopic brain tissues, even outside of the head region, demonstrating the instructive functional role of this large-scale voltage pattern for brain development ^88^.

Although the bioelectric prepattern that regulates brain morphology in this system is known, it is not understood how the *multicellular* patterns are collectively interpreted and executed across the field of cells in the neural plate and surrounding ectoderm. How is this tissue-level bioelectric pattern converted into specific cell-level decisions and genetic expression patterns across cells, leading to correct large-scale brain morphology? Here, we specifically address how large-scale bioelectric patterns in tissue interact with gene-regulatory circuits to establish the species-specific target morphology. We first developed a system in which machine learning could be deployed to search for empirically-testable, minimal, predictive models that integrate simplified bioelectric and transcriptional mechanisms. Then we used this methodology to produce and analyze a specific model of bioelectric control of *Xenopus* embryonic brain development. We combined *in silico* analysis and *in vivo* experimentation to elucidate the information-processing mechanism of the model and verify the consequent predictions. We show not only that the is model scale-invariant, but also that the spatial bioelectric pattern of the tissue controls gene expression via a simplified higher-order spatiotemporal information integration mechanism. We also show that there is functional specialization among cells, with cells at the voltage inflection points within the bioelectric pattern being important for correct pattern recognition and collective cellular decision making. Interestingly, we observe that the genes in a cell are responsive to the voltages of faraway cells. Furthermore, we found division of labor among the genes: genes are differentially sensitive to different aspects of the overall bioelectric pattern. This mechanism led to the counter-intuitive prediction of robustness, plasticity, and sensitivity of bioelectric patterns governing *Xenopus* embryonic brain development – predictions that we then confirmed *in vivo*. Taken together, this work demonstrates the tractability of a combined *in silico/in vivo* approach for identifying system-level features of a multi-modal morphogenetic process involving both biophysical and genetic components.

## Methods

### Model

We focus on neural development in *Xenopus* embryos at stage ~15 ^96–99^. We designed a minimal model of the neural plate tissue (Fig. 1F), characterized by the following biologically plausible features: a) a 2-dimensional lattice topology approximating the somewhat flat structure of the neural plate; b) topographic connectivity, realistically representing the connectivity only among neighboring cells in the tissue; c) symmetric connectivity of the tissue representing the homotypic gap junctions (GJ) conducting the flow of ions and an intercellular gene regulatory network (iGRN), conducting the flow of gene products, that assumes no directionality (e.g., PCP); d) an equivalent-circuit design of the bioelectric circuit consisting of just a pair of generic hyperpolarizing and depolarizing ion channels and voltage-gated gap junctions that was previously used to model the emergence and repair of the endogenous voltage pattern ^93^; e) a set of a minimum of six genes in each cell (representing the six genes linked to bioelectric circuits – *Notch, Otx2, Pax6, Emx, Xbf1/Foxg1, Sox2*) connected by a generic recurrent network (since the exact connectivity is not fully known we left it to machine learning to learn an appropriate one) and all cells containing the same GRN; f) a bidirectional connectivity between the cellular voltage and the genes of a single cell where the voltage acts as an “external input” to some genes, influencing their expression, and in turn the activity of a set of genes modulating the ion channel conductance (here again, since the exact nature of this relationship is not known we left it to machine learning to explore it); and g) various time constants associated with the gene activities and the connections from voltage to genes, genes to voltage, and within the iGRN, representing the heterogeneity of timescales associated with the respective modes of communication. The model was simulated using standard Euler integration for about 1000 steps starting from uniform initial conditions (details in SI. 2).

### Pattern discrimination problem

While we are interested in spatial computation in tissues (groups of cells) in general, we designed this model specifically to understand how *Xenopus* embryonic bioelectrical prepatterning leads to correct brain morphogenesis—to develop a normal brain. To that end, we set up a minimal version of this problem for our model to solve, which we will henceforth refer to as the *pattern discrimination problem*: recognizing the correct (endogenous) voltage pattern across cells by activating a set of genes and repressing them upon recognizing other patterns that are known to result in abnormal brain morphologies (thus recapitulating the results of *in vivo* experiments on role of bioelectric contrast prepatterns in *Xenopus* brain development ^76,88,89,91–93,100^). The input to the model is a voltage prepattern representing either the endogenous bioelectric contrast pattern (henceforth referred to as the “correct” pattern – Figure 1A-B) or its abnormal versions, namely fully depolarized or fully hyperpolarized (referred to as the “incorrect” patterns – Figure 1C-D). The output of the model is a gene expression pattern. The desired relationships between the inputs and outputs are as follows: 1) the correct voltage input pattern should result in the activation of the genes (positive activity); and 2) the incorrect voltage input patterns should result in the repression of those genes (negative activity).

### Machine learning search

Given the complex recurrent nature of this class of models and the necessity to begin to develop Artificial Intelligence tools that assist human scientists in discovering models of morphogenetic regulation, we used machine learning methods to search for an appropriate connectivity among its components and the associated parameters that enabled a model containing 24 cells (4×6 lattice) to solve the pattern discrimination problem (Fig. 1F). We used a combination of genetic algorithm (GA) and gradient descent (GD) to train the models. GA is analogous to biological evolution, whereas GD is analogous to learning during the lifetime of an animal. Our motivation for using this particular combination of search algorithms is not to emulate biological *learning* per se, but to discover an appropriate model connectivity and parameters that serve as a good, predictive model to explain the physiological and functional data on bioelectric control of brain patterning; the mixture of evolutionary and ontogenetic timescales in our searches is meant to be purely pragmatic. The overall search began with GA-enabled search for an appropriate network structure (including the numbers of nodes and edges), followed by GD to further fine-tune the parameters of the best network discovered by the GA, with the structure fixed. We used the standard microbial GA ^101^, which is based on horizontal gene transfer and a simple tournament selection process: two randomly chosen individuals from the population are pitted against each other and the ‘winner’ transmits randomly chosen portions of its genetic material to the ‘loser’ based on a specified “infection rate”. This process is repeated for thousands of iterations at the end of which the best performing individual (model) is chosen for further refinement using GD. We used a “resilient backpropagation” ^102^ flavor of GD for this purpose; this method relies only on the sign (not the magnitude) of the gradients for updating the parameters at every iteration. Our goal was to elucidate a set of conditions *sufficient* to solve the pattern discrimination problem, embodied by a single model. Hence, we chose a single high-performing model for subsequent analysis and examined the evolution of its ability to discriminate between correct and incorrect voltage patterns (SI. 3-5). Further details are in SI2-5.

### Performance measures

The quality of a model and its components was evaluated using the following performance measures (detailed definitions in SI. 3,4). The “discrimination error” of a gene is defined as the weighted mean distance between the observed spatial expression patterns and the target patterns corresponding to the endogenous, depolarized, and hyperpolarized input voltage patterns. The “discrimination score” of a gene is defined as the change in the observed discrimination error relative to the maximum possible error. The “performance score” of a model is defined as the inverse of the ratio between the mean observed and the maximum possible gene discrimination errors. Both the gene discrimination and the model performance scores range between 0 and 1, with the extrema corresponding to the negative and positive discriminations and the resulting model performance and a value of 0.5 indicating the expected discrimination and performance expected from a randomly parameterized model.

### Causal integration analysis

To investigate the causal relationship between voltage pattern and gene expression in our model we employed the framework of “multi-timescale causal influence” introduced in ref^103^. Specifically, the amount of first order causal influence (CI) exerted by the voltage pattern on gene expression is given by the following “Jacobian” tensor:

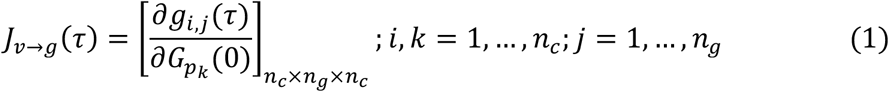

The tensor *J_v→g_*(*τ*) captures the sensitivity (positive or negative) of every gene in every cell, *g_i,j_*, at time τ with respect to the voltage of every cell, as determined by the conductance of the polarizing ion channel, *G_p_k__*, at time *t* = 0 for any *τ* ≥ 1. The larger the absolute value of an element of *J_v→g_*(*τ*), the more causally sensitive gene *g_i,j_* is at time *τ* to the initial voltage of cell *k*. The amount of second order CI exerted by the voltage on gene expression is given by the following “Hessian” tensor:

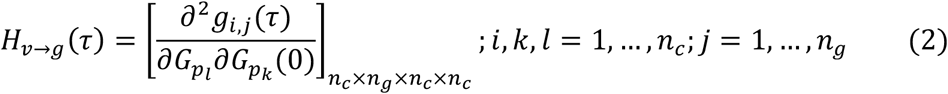

The tensor *H_v→g_*(*τ*) captures the second-order sensitivity (i.e., sensitivity of sensitivity, which can also be positive or negative) of every gene in every cell, *g_i,j_*, at time *τ* with respect to the voltages of every pair of cells, as determined by the conductance of the corresponding polarizing ion channels, (*G_p_l__, G_p_k__*), at time *t* = 0 for any *τ* ≥ 1. It can also be conceived as a measure of context-dependency – the extent to which the influence of a cell’s voltage on gene expression depends on the context provided by the voltage of another cell. Alternatively, it can be interpreted as the amount of collective influence exerted by the voltages of pairs of cells on the gene expression. The larger the absolute value of an element of *H_v→g_*(*τ*), the more causally sensitive gene *g_i,j_* is at time *τ* to the initial voltages of the pair of cells *l* and *k*. Here we used the conductance of the polarizing ion channel, *G_p_*, of a cell as a surrogate for its voltage since the former is fixed throughout the simulation and fully determines the voltage profile of tissue ^93^ (Fig 5A).

### Animal Husbandry

*Xenopus laevis* embryos were fertilized *in vitro* according to standard protocols in 0.1X Marc’s Modified Ringer’s solution (MMR: 10mM Na^+^. 0.2 mM K^+^, 10.5 mM Cl^-^, 0.2 mM Ca^2+^, pH 7.8;) ^104^. Embryos were housed at 14-18°C and staged according to Nieuwkoop and Faber ^96^. All experiments were approved by the Tufts University Animal Research Committee (M2020-35) following the guide for the care and use of laboratory animals.

### Microinjections

Capped synthetic mRNAs generated using the mMessage mMachine kit (Ambion) were dissolved in nuclease-free water and injected into embryos immersed in 3% Ficoll using standard protocols ^104^. Each injection delivered ~ 1-2 ng of mRNA (per blastomere) into the middle of a cell in the animal hemisphere of embryos at the four-cell stage. Constructs used were: *Kv1.5* ^105^, *dominant-negative Kir6.1 pore mutant – DN-Kir6.1p* ^31^, and *β-galactosidase*.

### Membrane voltage imaging with DiBAC_4_(3):CC2-DMPE

DiBAC_4_(3) and CC2-DMPE ratiometric reporter dyes (Invitrogen) were used as per the standard protocol ^106^. Briefly, CC2-DMPE stock (5 mM) was dissolved 1:1000 in 0.1X MMR and the embryos were incubated in the dark for 1 hour followed by five washes with 0.1X MMR. DiBAC_4_(3) stock (1.9 mM) was dissolved 1:1000 in 0.1X MMR and the CC2-DMPE stained embryos were then incubated in the dark in this solution for 30 min followed by five washes with 0.1X MMR, and then visualized under an Olympus BX-61 microscope equipped with a Hamamatsu ORCA AG CCD camera controlled by MetaMorph software (Molecular Devices).

### β-Galactosidase Enzymatic Detection

Tadpoles were fixed at stage 45 for 30 mins in MEMFA at room temperature, washed twice in PBS with 2 mM MgCl_2_, and stained with X-gal (Roche Applied Science, Indianapolis, IN, USA) staining solution at 37°C for at least 3 hours. Tadpoles were then rinsed three times in PBS followed by dehydration through sequential incubation in 25%, 50%, 75% and 100% methanol. Tadpoles were then incubated in 30% H_2_O_2_ in methanol overnight for bleaching, washed in 100% methanol, and sequentially (100%, 75%, 50%, 25%) rehydrated to PBS and imaged.

### Statistics

Statistical analyses were performed using GraphPad Prism. At least three independent experiments were conducted with N>50 embryos for each treatment group, using embryos collected from multiple animals across independent clutches. Data were analyzed by ANOVA (with Tukey’s multiple comparison post-test).

### Data Availability

All data generated or analyzed during this study are included in this article and are available from the corresponding author upon request.

## Results

### 1. Machine learning discovered a model that solves the voltage pattern recognition problem

Given the complex recurrent nature of this class of models and the necessity to begin to develop Artificial Intelligence tools that assist human scientists in discovering models of morphogenetic regulation, we used machine learning methods to search for a connectivity among model components and the associated parameters that enabled solution of the pattern discrimination problem (for details, see Methods and SI. 2-5). Our goal was to elucidate a set of conditions *sufficient* to solve the pattern discrimination problem, embodied by a single model. Hence, we chose a single high-performing model for subsequent analysis and examined the evolution of its ability to discriminate between correct and incorrect voltage patterns (SI. 3-5).

The design of the high-performing model we analyzed further, comprising the connectivity and the associated parameters, is depicted in Fig. 3 as a simplified two-cell version of the full model (Tables S1–S8) comprising about 20% of the strongest connections (10% of the most positive and 10% of the most negative). As can be expected, the strongest connections (thick edges) of the model are those that run from the voltage to a few genes so that the input could be reliably transmitted to the genes, and those connecting a small subset of the genes both within a cell and between cells so that the transmitted message could be reliably processed and communicated to other cells. We next analyze in detail the behavior and dynamics of this model.

**Figure 3.**
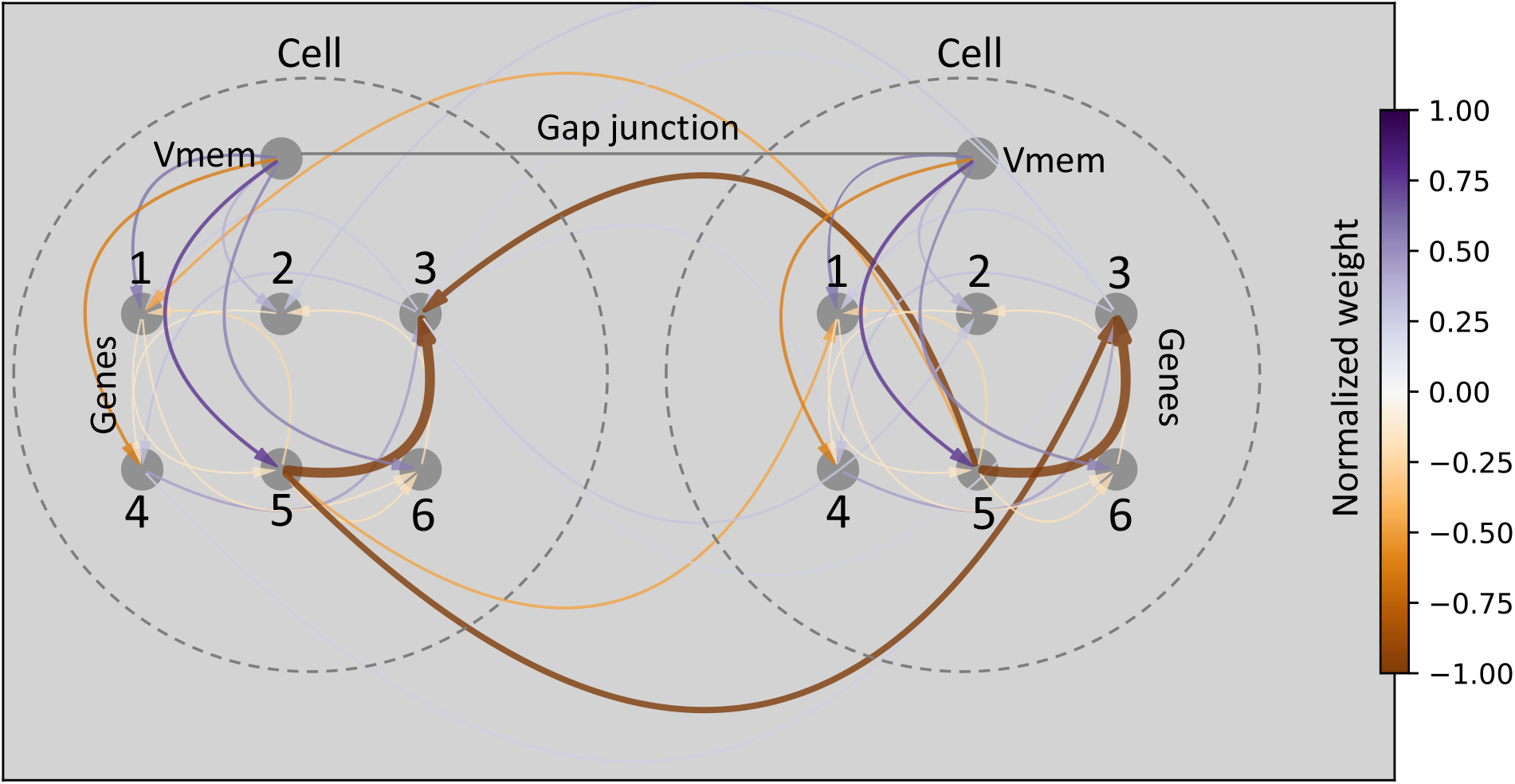
Architecture of the chosen high-performing model. This depicts the connectivity of a two-cell version of the full model; the rest of the network is symmetrical to this version (full model in SI. 1). For clarity, we show only about 20% of the strongest connections (10% of the most positive in purple and 10% of the most negative in orange/brown); edge weights are linearly normalized to the range (−1,1). The most prominent connections run from the voltage to gene 5 (purple), with a positive weight, and among genes 3 and 5 both within a cell and between cells (brown), with a negative weight. Connections running from genes to voltage do exist in the full model, but they were filtered out due to their weak weights. The gap junction is depicted for visual purpose only; its weight is not set to scale since it is dynamically voltage-modulated.

On average, genes in the model successfully discriminate between the correct and incorrect voltage patterns: when averaged over all cells and all genes, expression is increased for the correct voltage pattern and decreased for the incorrect pattern (Fig. 4A). Average magnitudes of expression are relatively low, about +0.5 for activation and −0.2 for repression, compared to the ideal values of +1.0 and −1.0 respectively, yielding a suboptimal performance score of 0.64, but they are significantly higher than expected for a randomly parameterized model (SI.5). Importantly though, the gene expressions for the correct and incorrect voltage patterns clearly diverge over the time course. When we look at individual genes, this separation is largest for gene 6, as indicated by the corresponding discrimination score (Fig. 4A; bottom right inset). For this reason, we term gene 6 the “discriminator gene” and focus on its behavior in the analyses below.

**Figure 4.**
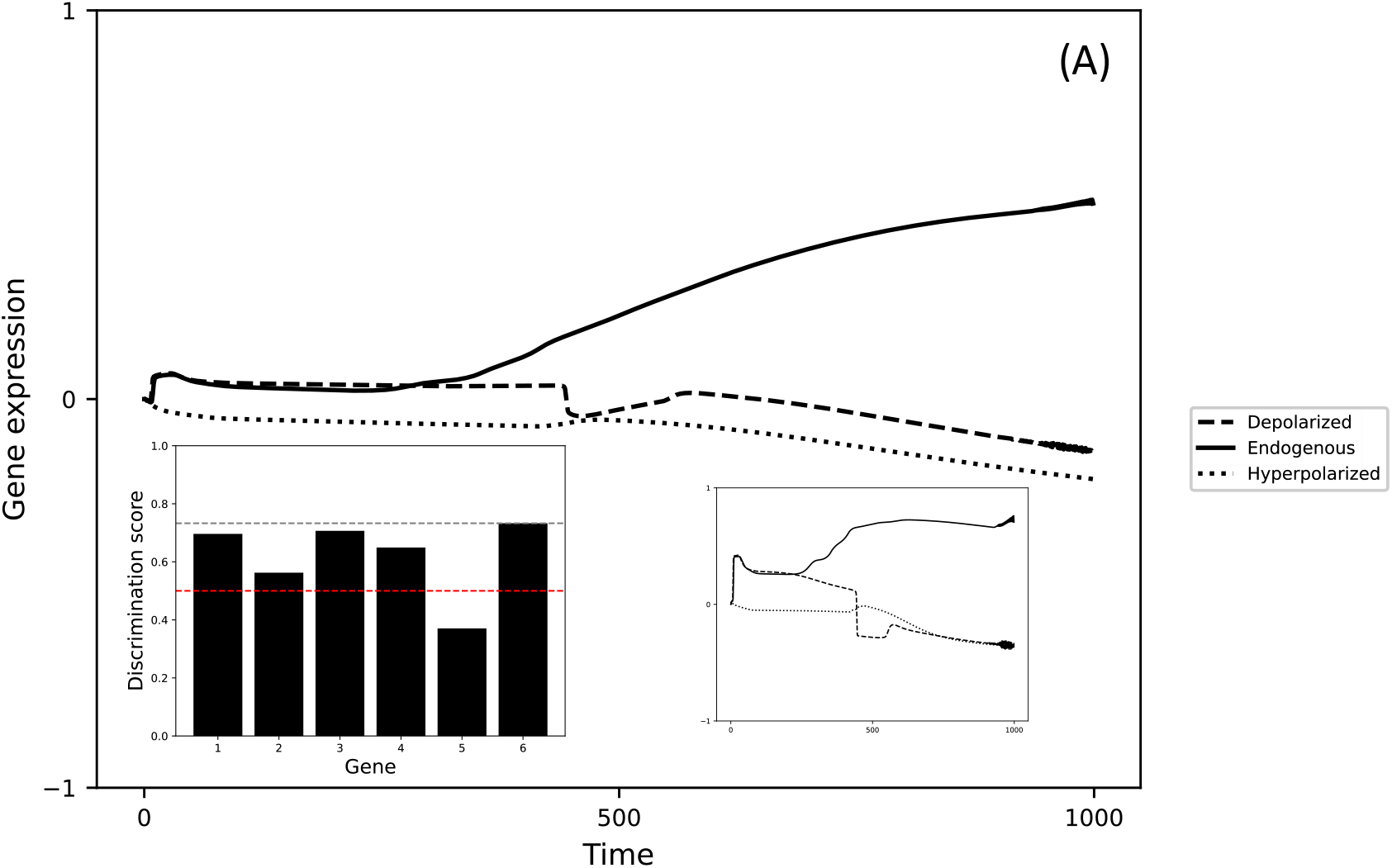
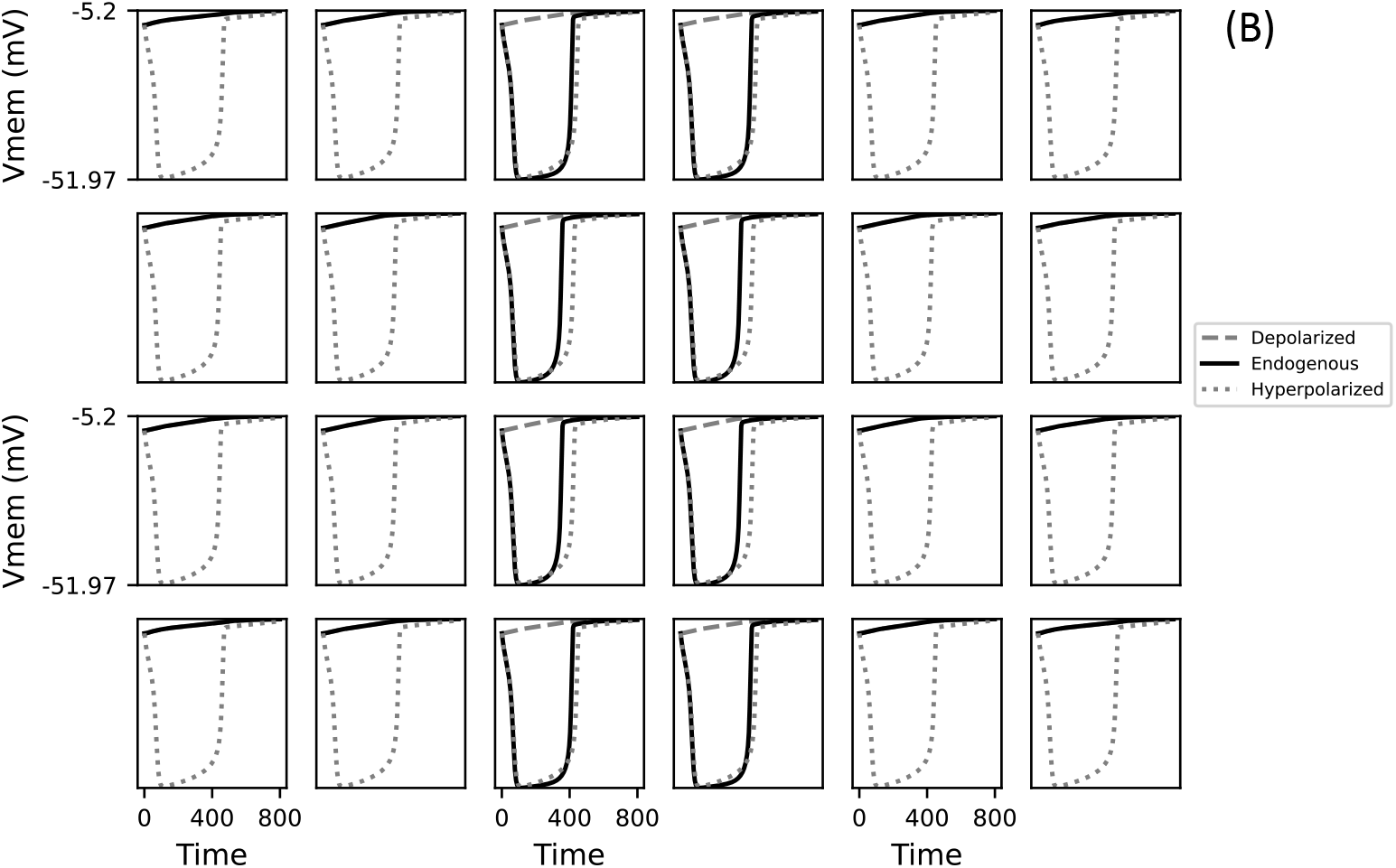
The best fit model recapitulates the relationship between voltage pattern and gene expression as well as discriminates between the correct and the incorrect voltage patterns. Each square in the 4×6 layout of the tissue refers to an individual cell. (A) Timeseries of the mean normalized gene expression, averaged over all cells and genes, for the three different input conditions depicted in Figure 1B-D: “Depolarized” (all cells are initially depolarized), “Endogenous” (the left and right two columns are initially depolarized with the middle two columns hyperpolarized), or “Hyperpolarized” (all cells are initially hyperpolarized). Insets: (Left) depicts the discrimination scores of the individual genes. The red dashed horizontal line indicates the expected discrimination score from a randomly parametrized model; the black dashed horizontal line indicates the value for gene 6, which had the highest score. (Right) depicts the gene expression timeseries for gene 6. B) Timeseries of Vmem for each cell in the model under each of the three input conditions.

Interestingly, the temporal dynamics of the voltage pattern in this model also recapitulate the developmental appearance of the bioelectric pattern *in vivo* even though the model was not specifically trained to develop it (solid black lines in Fig. 4B). At the beginning, our model exhibits a flat depolarized pattern across all cells; then, fairly early in the simulation, the cells in the center (modeling the neural plate) hyperpolarize to produce the characteristic voltage contrast seen *in vivo*, before reverting to the flat depolarized pattern about halfway through the simulation. This corresponds to what we see in this region between developmental stages 14 and 20 *in vivo* ^88^. These observations suggested that even though the quantitative performance of the model is not ideal, its quality warranted further analysis.

### 2. Model solves the pattern discrimination problem even in larger tissues despite not having been selected for such capability: scale robustness

One central aspect of biological control mechanisms is that they often have properties that were not specifically selected for, by virtue of generic (inherent) properties of networks ^8,107,108^. We were interested to uncover any novel emergent features in our model, focusing especially on size control. The ability to establish correct overall pattern despite differences in tissue size or available cell number ^109–113^, a key competency of developmental and regenerative morphogenesis, is still poorly understood in models based on reaction-diffusion networks ^54,114,115^ due to their limited scaling robustness.

Interestingly, our model successfully solved the pattern discrimination problem even in larger ‘tissues’ (180 cells compared to the original 24 cells), even though it was not specifically trained for that purpose (Fig. 5). Similar behavior is observed with even larger tissues (Fig. S12). This suggests that the learned dynamics are general and were not specific to a particular tissue size or shape features of the training. Moreover, this means that insights we derive from the analysis of the model should be generally applicable to many tissue sizes. We’ve previously hypothesized that this ability to rescale could be attributed to particular model parameters: it has a homogeneous connectivity and all the cells have the same GRN ^103^. In addition, the qualitative behavior of the discriminator gene is also preserved – its activity in the flanking columns is relatively lower compared to the columns closer to the center (Fig. S2), suggesting that the dynamics of the original model have scaled up (Fig. S3).

**Figure 5.**
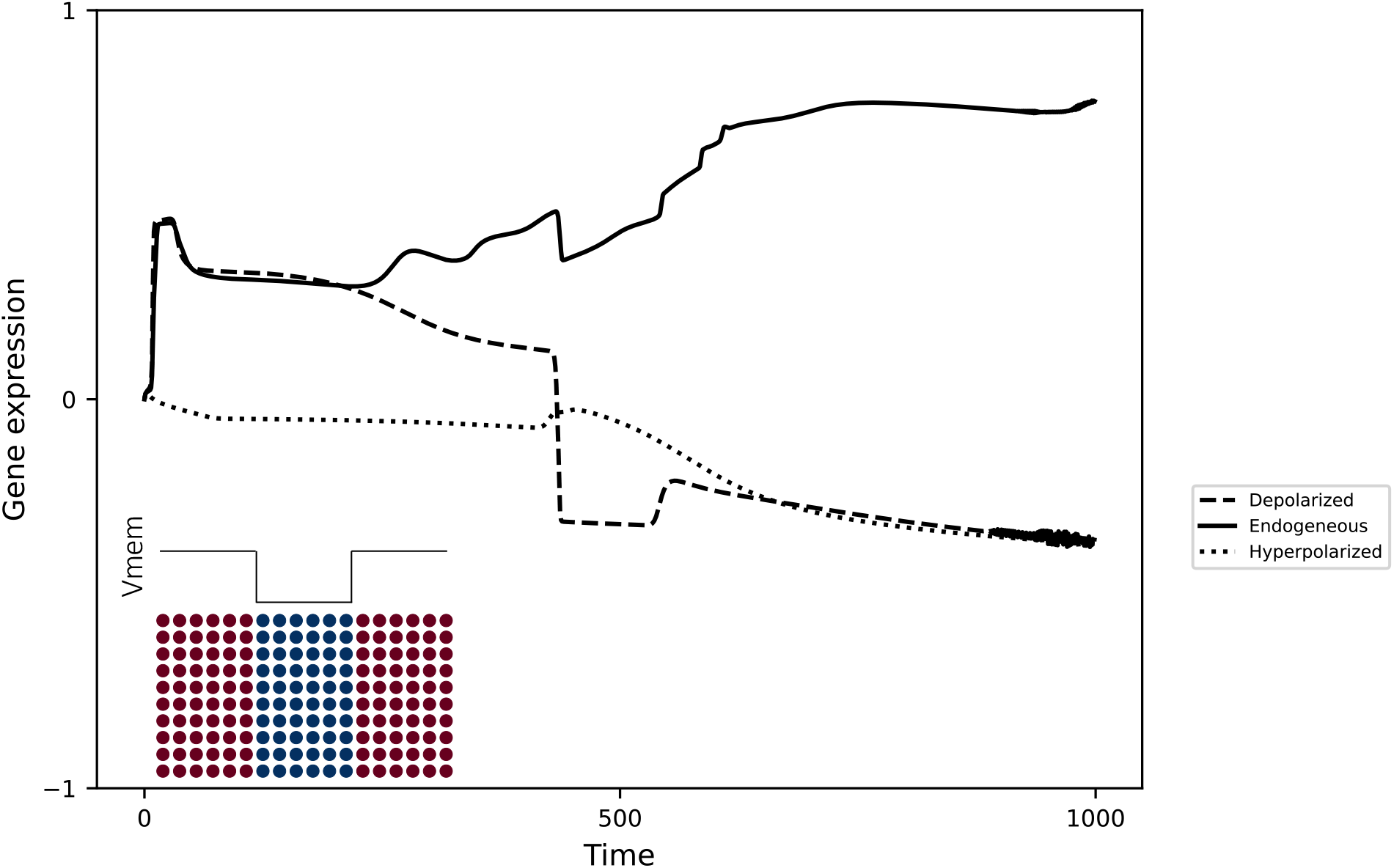
The best fit model is scale invariant (larger tissues) when solving the voltage pattern discrimination problem. The timeseries of the mean expression, averaged over all cells, of the discriminator gene in a scaled up 10×18 model when initialized with the endogenous voltage pattern (inset; left and right six columns are depolarized, and the middle six columns are hyperpolarized), showing that it successfully discriminates between the correct and the incorrect voltage patterns.

### 3. Deciphering high-level information-processing mechanisms employed to solve pattern discrimination problem

We next performed a detailed analysis of the model to decipher some of the high-level information-processing mechanisms and organizational principles it employs to solve the pattern discrimination problem. In the following, we specifically asked: a) Does functional specialization exist among the components (cells, genes, modules, etc.)? b) Does a division of labor exist in terms of the scales (distributed or local) at which information-processing is performed? c) Are there high-level relationships among the components that facilitate pattern discrimination? d) How is the voltage pattern stored and processed in the tissue?

#### a. Cells surrounding voltage transition points are more important for recognizing the bioelectric contrast pattern

To determine whether different cells in the tissue play different roles in the collective recognition of the endogenous voltage pattern, we performed an *in silico* cell “knockout” analysis. Specific cells were frozen by clamping the states of the corresponding genes at zero and dropping their biases to negative infinity, thereby rendering them ineffectual both as inputs and output, and the performance of the model was then computed.

Knocking out the cells surrounding the voltage transition points of the endogenous pattern – at the boundaries of the central band where there are large changes in the spatial profile of the voltage pattern – resulted in a greater drop in performance (Fig. 6). In other words, cells closer to the voltage transition points are more important in recognizing the pattern compared to the cells further away. Specifically, the four cells at the center of the 4×6 tissue are the most significant contributors to performance (Fig. 6A). By virtue of scale robustness, this property also extends to the larger 10×18 tissue where small 2×3 patches of cells were knocked out. There again, the cells flanking the center of the tissue are the most important (Fig. 6B). Though there are slight differences between the smaller and the larger tissues with regard to the exact spatial regions that are most important, the general principle holds: cells coinciding with changes in the pattern contribute more to its discrimination. A potential explanation for these observations is that the voltage transition points comprise the only difference between the endogenous pattern and the uniform patterns, suggesting that this strategy may computationally optimal.

**Figure 6.**
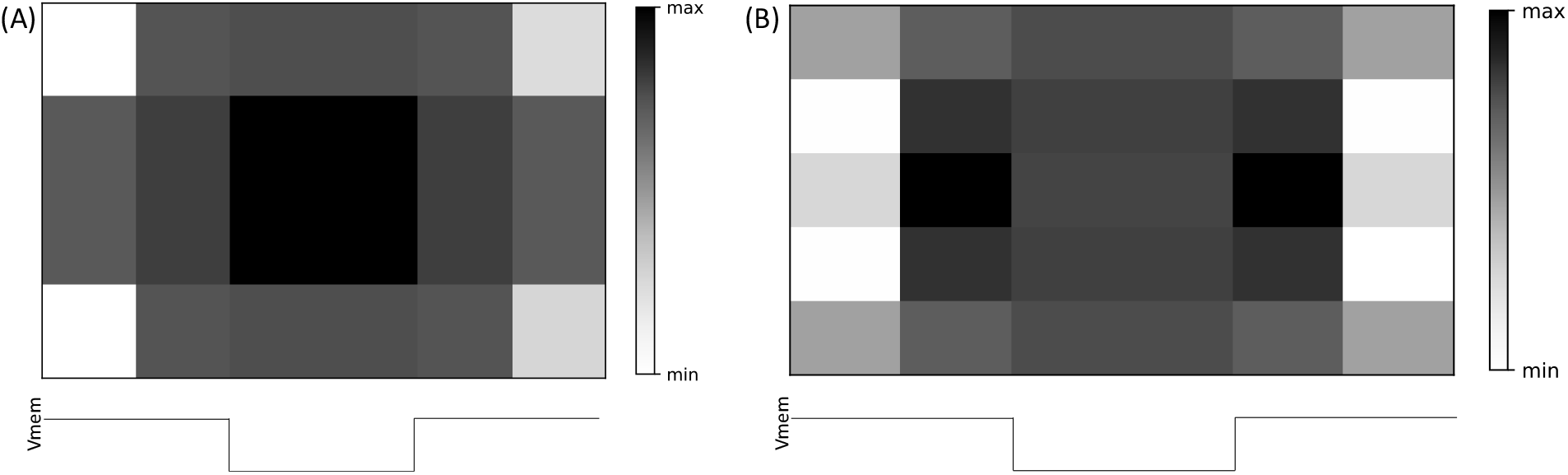
A spatial map illustrating the drop in model performance due to knocking out individual or patches of cells in the tissue. A cell or a patch was “knocked out” by freezing the states of its corresponding genes at zero and dropping their biases to negative infinity, thereby rendering them ineffectual both as inputs and outputs. (**A**) A 4×6 tissue where individual cells were knocked out. (**B**) A 10×18 tissue where 2×3 patches of cells (each block is a patch) were knocked out. The darker the shade of a block, the higher the drop in performance due to knocking out that block.

#### b. A simplified higher-order control mechanism characterizes the relationship between the endogenous voltage pattern and gene expression

To characterize the high-level relationships between the endogenous voltage pattern and tissue-level gene expression, we attempted to express the dynamic behavior of the discriminator gene in terms of its dynamic sensitivities to voltage captured in the time-dependent Jacobian and Hessian tensors (Eqns. 1 and 2). Due to the 4-fold reflection symmetry of the 2D lattice-structured tissue (Fig. 1F), we computed the sum of sensitivities of the discriminator gene, corresponding to the endogenous input pattern, only for the cells in the top left quadrant of the tissue (details in SI. 12). The extent to which these quantities can reconstruct the dynamic timeseries of gene expression reflects the amount of control that voltage exerts over gene expression; a simple example of this is described in SI 9.

We found that the dynamic profile of the discriminator gene expression corresponding to the endogenous input pattern could be almost fully recovered with just the Hessian (Fig. 7), with a match significantly higher than random expectation (Fig. S8). On the other hand, the Jacobian alone produced a reconstruction with a lower quality of match that was not significantly different from random expectation (Fig. S8). Taken together, these observations suggest that the voltage pattern exerts control over the discriminator gene expression almost exclusively at the second-order level. In other words, it is the voltages of pairs of cells, rather than of single cells, that influences gene expression. A possible explanation is that pattern discrimination requires a comparison of voltage of different cells, which the Hessian presumably aids by containing the differential information. The above observations also suggest that the voltage pattern controls tissue-level gene expression in an effectively feed-forward manner (SI 9) despite the complex recurrent connectivity between them (Figure 4; also demonstrated by the fact that severing the feedback connections from the genes to the voltage partially affects the pattern discrimination ability of the model, as shown in Fig. S13). In other words, the bioelectric pattern exerts a “canalized” control over the discriminator gene’s expression by effectively linearizing the complex nonlinearity of the network – an observation that cannot be discerned from an inspection of the static connectivity of the tissue (Fig. 3) alone; a simple analog of this is described in SI 9.

**Figure 7.**
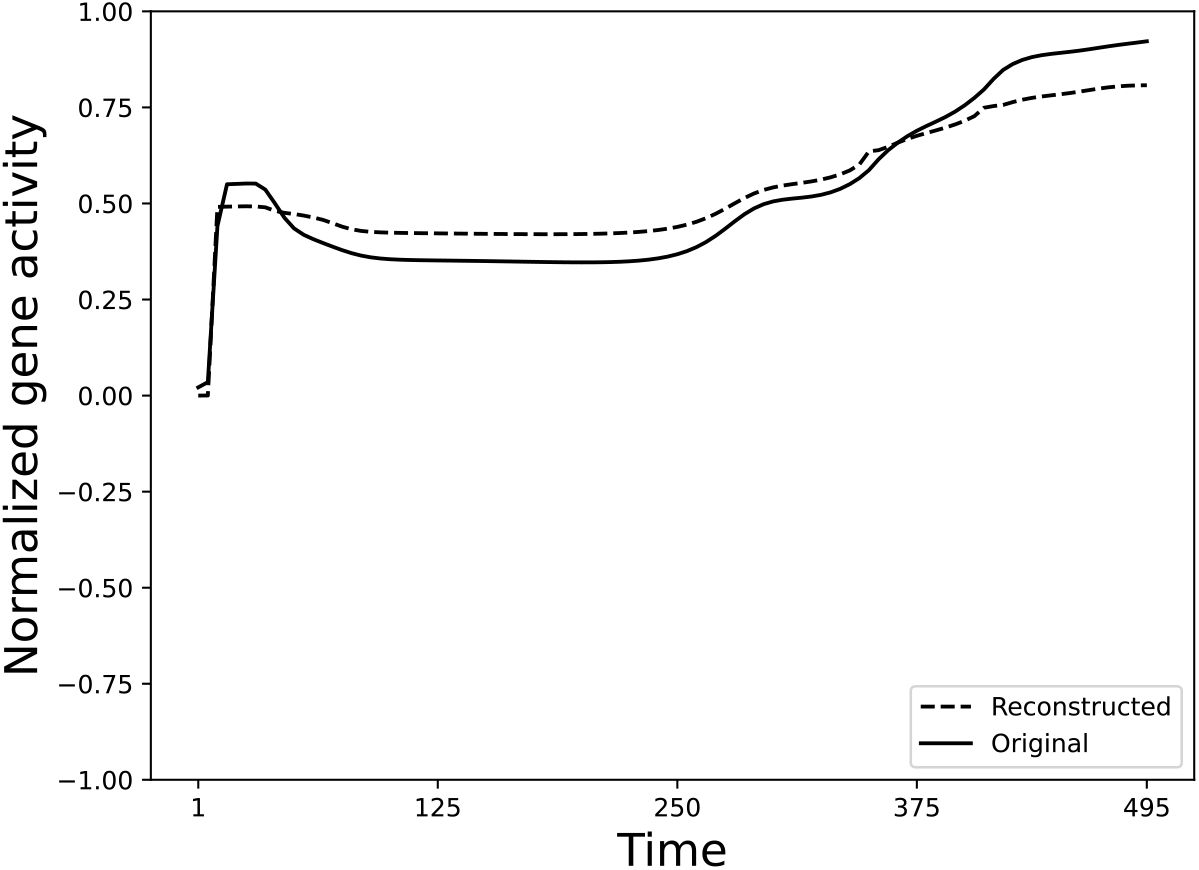
A comparison between the normalized original gene expression timeseries of the discriminator gene (dashed) for the endogenous input pattern and the corresponding reconstruction (solid) obtained from the second-order (Hessian) influence of the endogenous voltage pattern. Due to the model’s symmetry, only the genes contained by the cells in the 2×3 top left quadrant of the full 4×6 tissue (dashed boxes in Fig 9) were considered for these calculations; the Hessian depicted here is the sum of all Hessians corresponding to each cell in the quadrant.

#### c. Long-distance influence and a scanning-like mechanism characterize bioelectric control

To further characterize the information-processing mechanisms underlying the bioelectric control of tissue-level gene expression, we analyzed the spatiotemporal organization of the Hessian networks underlying the reconstruction of the discriminator gene expression timeseries shown in Fig. 7 (details in SI. 12). We found that these networks are characterized by symmetry-breaking and long-distance influence, where the resemblance of the network with the original lattice structure (Fig. 1F) declines during the simulation, and some of the influencing cells tend to be distant (Fig. 8). This is somewhat surprising since the symmetric and topographic nature of the underlying connectivity model would suggest that the influences remain local. Moreover, the influences are dynamic and oscillatory in that they start out locally during the early time points (t=5), then switch to the other bands (t=50-410) and then back to being more local (t=495). During this process, the signs of the influence also change; for example, the dependencies on both the right band and the left band are negative at t=50 but switch to being positive at t=200. This oscillatory scanning-like behavior suggests that information is integrated over both space and time before a decision is made – a hypothesis that’s further supported by the observation that a similar mechanism is exhibited by the Jacobian as well (Fig. S10). A possible explanation is that since the genes in a cell have immediate access only to the cell’s own voltage but not that of other cells, an information integration mechanism that collects and supplies that information from across the tissue, such as the above, may be required.

**Figure 8.**
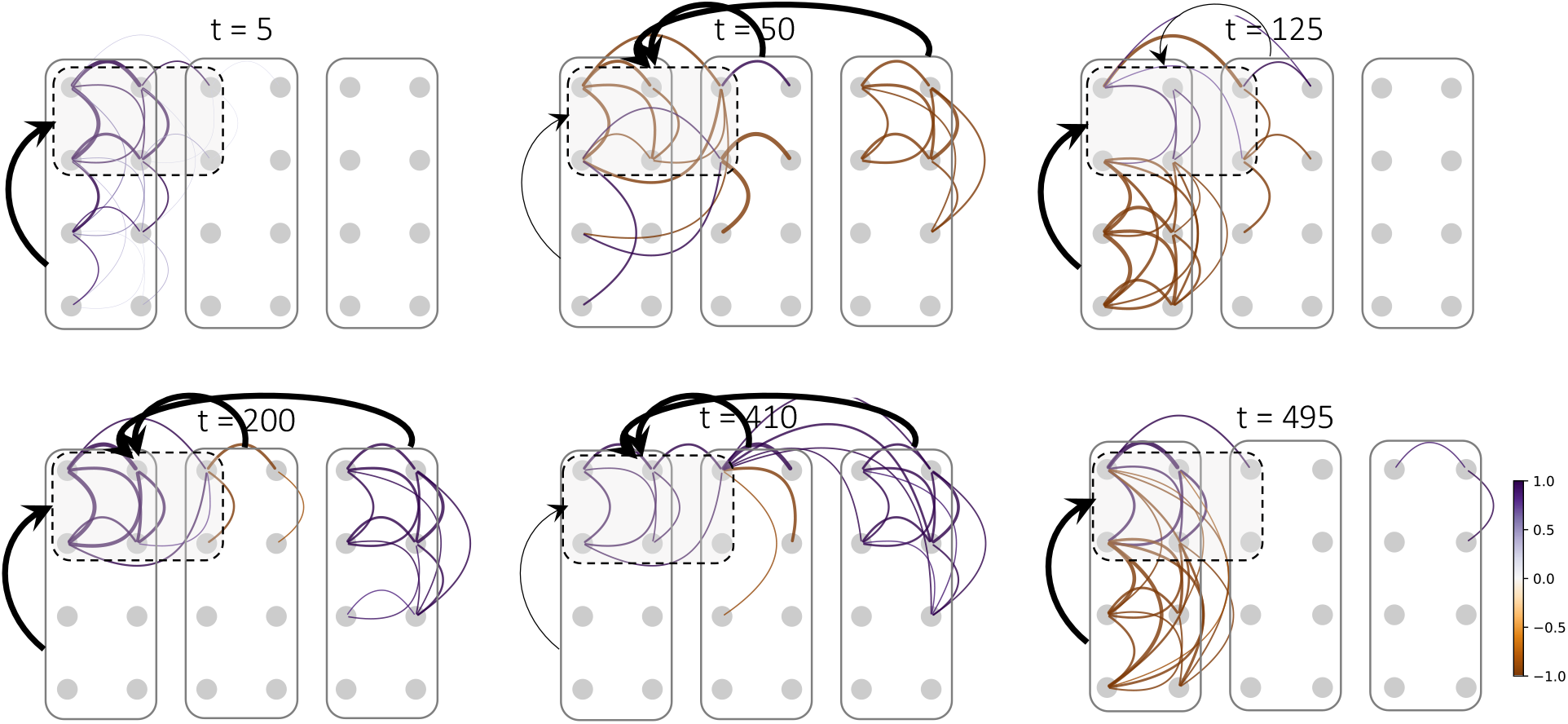
The spatiotemporal dynamics of the second-order control of gene expression by the endogenous voltage pattern. Each edge represents a pair of cells whose voltages collectively influence the expression of an average discriminator gene in the top left quadrant. The undirected edges indicate the influence exerted by the voltages of the corresponding pairs of cells on the discriminator gene; each edge is a visual representation of a single mean second-order derivative, averaged over both orderings of the pair. Due to the symmetry of the model, only the cells in the 2×3 top left quadrant of the full 4×6 tissue (dashed boxes) were considered for these calculations; the Hessian depicted here is the sum of all Hessians corresponding to each cell in the quadrant. Only about 10% of the observed edges is shown for clarity. Colors represent the strength and the sign of the dependency (strong purple means strongly positive and strong orange/brown means strongly negative).

#### d. Different genes are attuned to different spatial scales of the tissue: division of labor

We next analyzed the scales of sensitivity of each gene by computing the difference between the Jacobian (first-order) sensitivity to the whole tissue and that of the single cell that contains it, averaged over the cells in the top left quadrant of the tissue (details in SI. 12). We found that there is a unique apportioning of the gene sensitivities among the various spatial scales of the voltage pattern in that some genes (2 and 5) are more attuned to the pattern at the tissue level, some (genes 3 and 4) to the single-cell level, with others (genes 1 and 6) display a fine balance between the two scales (Fig. 9). This distribution of labor suggests that different genes play different functional roles tied to their respective spatial sensitivities. Indeed, we found that knocking out genes 2 or 5 affected detection of the non-endogenous uniform patterns but not the detection of the endogenous voltage pattern, while knocking out genes 4 or 6 affected detection of the latter (SI. 13). This implies that genes 2 and 5 are specialized for the homogeneous (incorrect) patterns, while genes 4 and 6 are specialized for the heterogeneous patterns. A possible explanation that ties these observations to the genes’ spatial sensitivity scales is that detection of homogeneity requires a tissue-level sensitivity, but heterogeneity requires a relatively smaller scale of sensitivity in this model. It appears that genes 2 and 5 sense whether the voltages across all cells in tissue are equal, whereas the other genes are sensing local differences in voltages. The tissue is deemed heterogenous if a local patch is heterogenous, whereas the tissue is homogenous only if all local patches are homogeneous, hence the differences in the corresponding scales of sensitivities.

**Figure 9.**
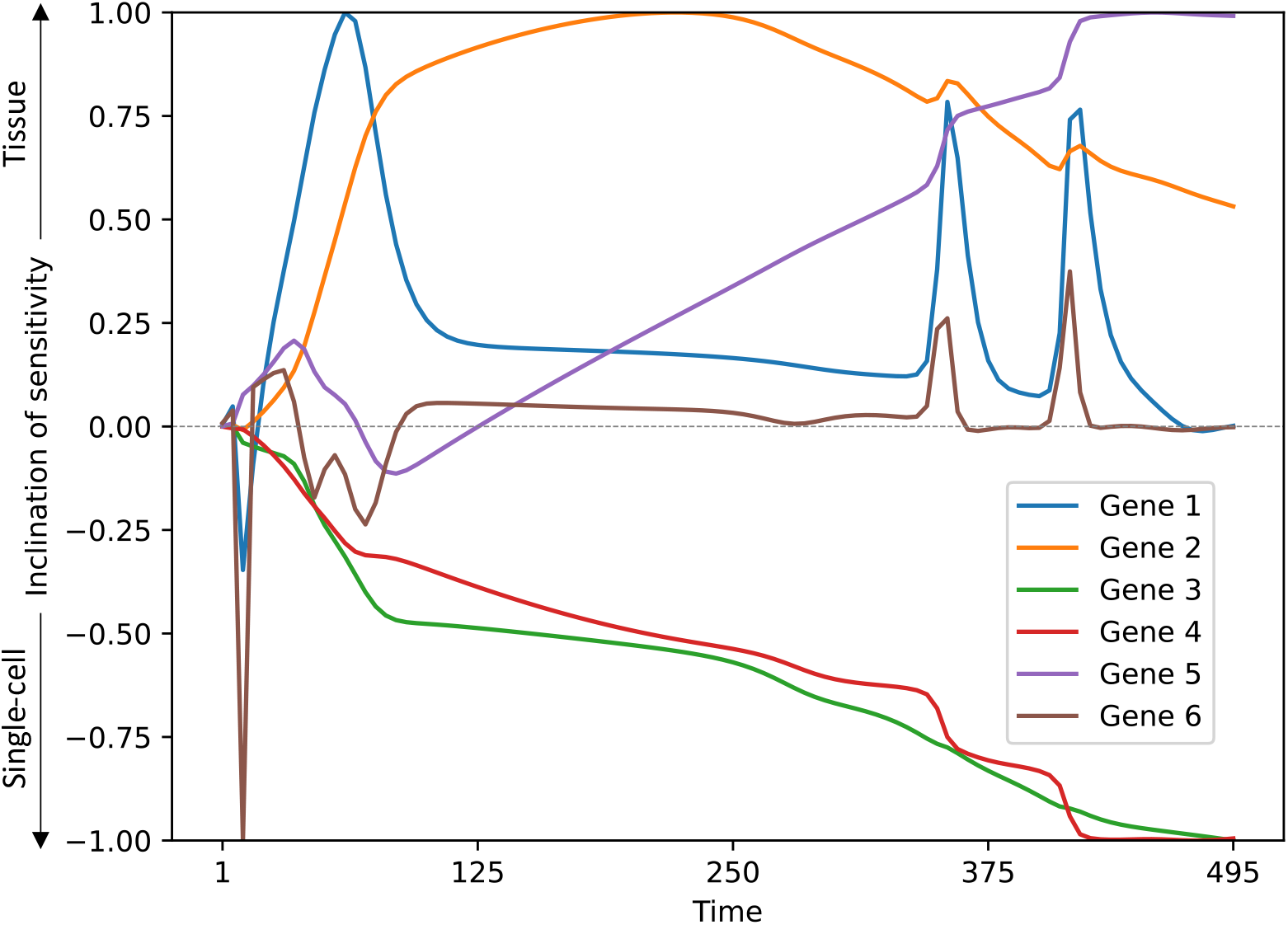
A division of labor exists among the genes in terms of their sensitivities to scales of voltage patterns. The difference between the first order (Jacobian) directional derivatives of the gene activities computed with respect to the voltage at the tissue level and the single-cell level tends to be positive for genes 2 and 5, negative for genes 3 and 4, and mostly balanced but interspersed with positive spurts for genes 1 and 6. The corresponding inclinations of sensitivity are towards the tissue level, single-cell level, and an intermediate level respectively. Due to the symmetry of the model, only the genes contained by the cells in the 2×3 top left quadrant of the full 4×6 tissue were considered for these calculations.

### 4. Model predicts robustness of voltage patterns in governing developmental brain morphology

We already know that incorrect voltage patterns can disrupt gene expression and morphogenesis ^28,29,88,91,93^ (Figs. 1B-D in Ref ^88^). We next asked whether our model can provide insight into the robustness of this system: which changes in the voltage pattern do not interfere with correct brain morphological patterning? Are there conditions where the overall shape of the voltage pattern is preserved but modifying certain aspects does disrupt normal brain morphological patterning? To that end we sought to test patterns that are: 1) suggested by some of the insights gained into the information-processing mechanisms above; and 2) experimentally tractable.

The analyses detailed in the previous section suggest that an asymmetric context-dependency exists among the cells: the extent to which the voltage of a cell influences the gene expression in other cells depends on the type of polarization of the other cells. Specifically, there’s a greater dependency on depolarized cells compared to hyperpolarized cells, on average, as seen in Fig. 8 and quantified in Fig. S5. In other words, depolarized cells exert a greater influence than hyperpolarized cells on the gene expression of other cells regardless of their polarization. This suggests that the proportion of depolarized to hyperpolarized cells in the tissue may be one of the factors that determines how the voltage pattern is processed. Consequently, we tested two types of modified endogenous patterns, namely, a “half-and-half” pattern and a “sharpened” pattern. In the half-and-half pattern, one half of the tissue is hyperpolarized and the other half is depolarized (in essence a step function pattern), whereas in the sharpened pattern the overall voltage pattern is maintained but fewer cells in the central band are hyperpolarized compared to the normal pattern.

The model predicted that the half-and-half pattern would be equivalent to the normal pattern, resulting in normal gene expression and normal brain morphology (Fig. 10A). Contrarily, it also predicted that the sharpened pattern would disrupt gene expression in the flanking columns (Fig. 10B), resulting in the development of a partially defective brain. We then tested these model predictions *in vivo*.

**Figure 10.**
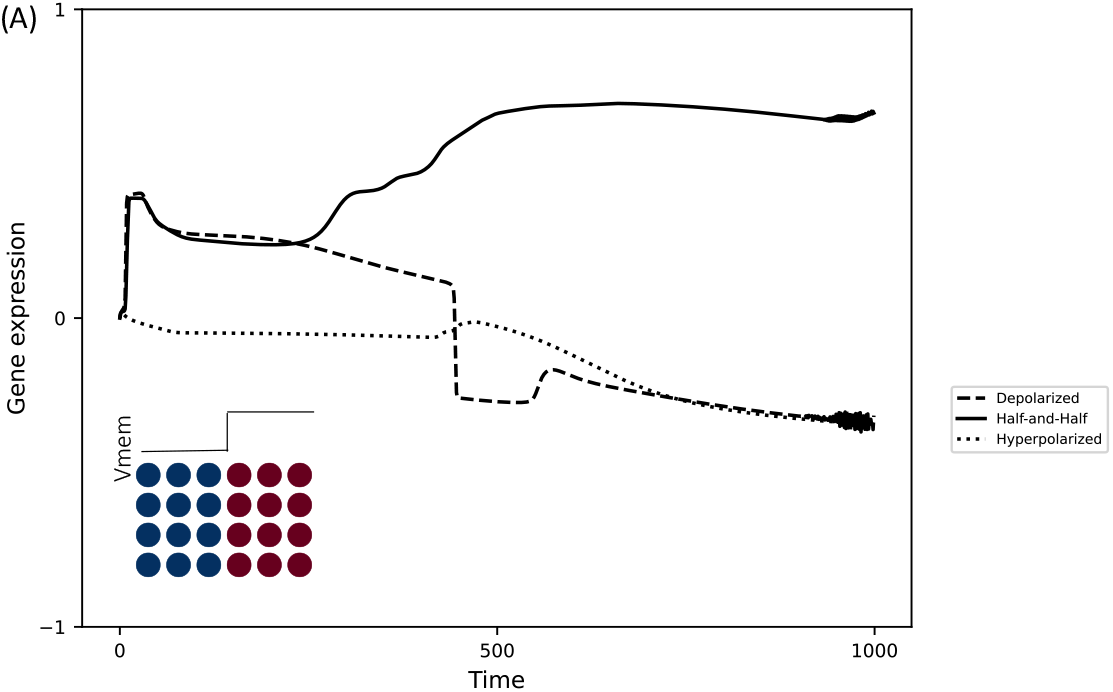
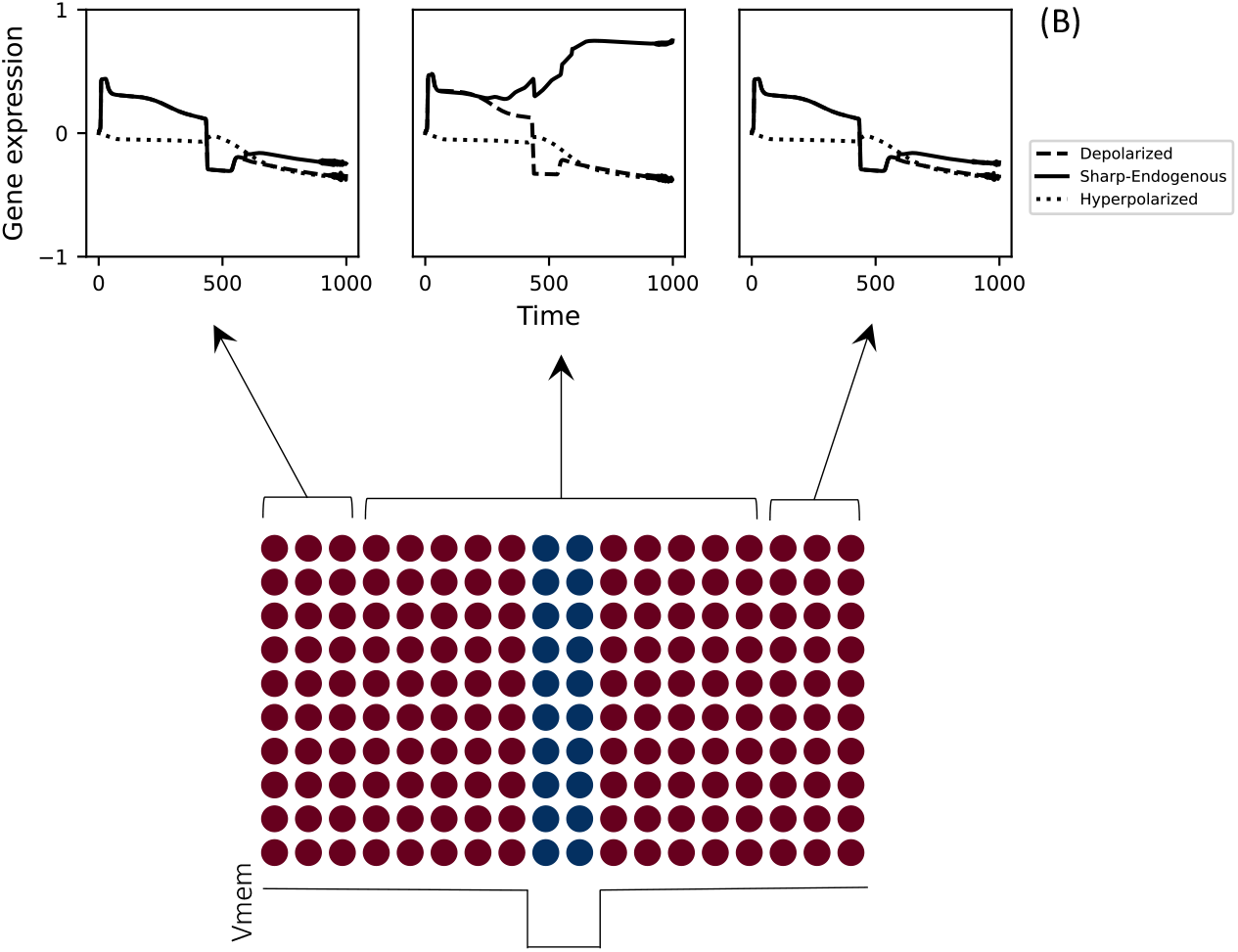
Model predictions about the robustness of the system. (**A**) A half-and-half (step function) voltage pattern where the left half is hyperpolarized, and the right half is depolarized (inset) should result in normal brain development as indicated by the mean gene expression of the discriminator gene, averaged over all cells, discriminating it from the other patterns. (**B**) A sharpened voltage pattern where the overall pattern is maintained but the hyperpolarized neural plate (central band) is only a third of its original normal size (bottom panel: only the middle two columns of the 10×18 tissue are depolarized) would result in an intermediate/non-normal brain development due to the failure of the mean gene expression to discriminate between the endogenous and the incorrect voltage patterns in the leftmost and the rightmost three columns (indicated by the arrows).

### 5. Empirical testing of prediction one: altering the voltage pattern regulating brain development in only one half of *Xenopus* embryo does not disrupt endogenous brain development

As described above, our previous work showed that disrupting the normal membrane voltage pattern in the neural plate and surrounding ectoderm resulted in significant abnormalities in large-scale brain morphology and function ^88,89,91–94^ (Fig. 1C-D). Hence our model’s first prediction (Figure 10A), that inducing an abnormal step function bioelectric pattern will *not* disrupt normal brain development, is counter-expectation. To test this prediction, we used a well-established strategy for altering voltage patterns in *Xenopus* embryos by overexpressing ion channels via mRNA microinjection ^31,40,88,91–93,116,117^.

The neural plate and surrounding ectoderm are largely derived from the two dorsal blastomeres of the four-celled *Xenopus* embryo ^118^ (Fig. 11A). To convert the endogenous voltage contrast pattern to a step function pattern, we co-microinjected *Kv1.5* (a voltage-gated potassium channel that hyperpolarizes ^105^) with lineage-tracer *β-galactosidase* mRNA into a single dorsal blastomere (Fig. 11A-C). Uninjected embryos and embryos injected only with *β-galactosidase* mRNA served as controls. Imaging of ~stage 15 embryos using voltage reporter dyes confirmed that *Kv1.5 + β-galactosidase* mRNA microinjection hyperpolarized only the injected half of the neural plate and surrounding ectoderm, changing the normal v-shaped voltage pattern seen in control embryos to a step function pattern (Fig. 11B,C). We then evaluated brain morphology and confirmed our microinjection tissue targeting at stage 45. At stage 45, control tadpoles showed normal brain patterning ^81^ (Fig. 11D &H). Surprisingly, as predicted by the model, the *Kv1.5 + β-galactosidase* mRNA-injected tadpoles also exhibited normal brain patterning (Fig. 11E & H). To demonstrate that this counter-expectation result is not specific to Kv1.5 channels but is due to the altered voltage pattern, we repeated the experiment with a different channel, Kir4.1, which is also known to hyperpolarize cells ^33,119^. Microinjecting *Kir4.1* mRNA in the same protocol also resulted in stage 45 tadpoles with normal eye and brain patterning (Fig. 11H).

**Figure 11:**
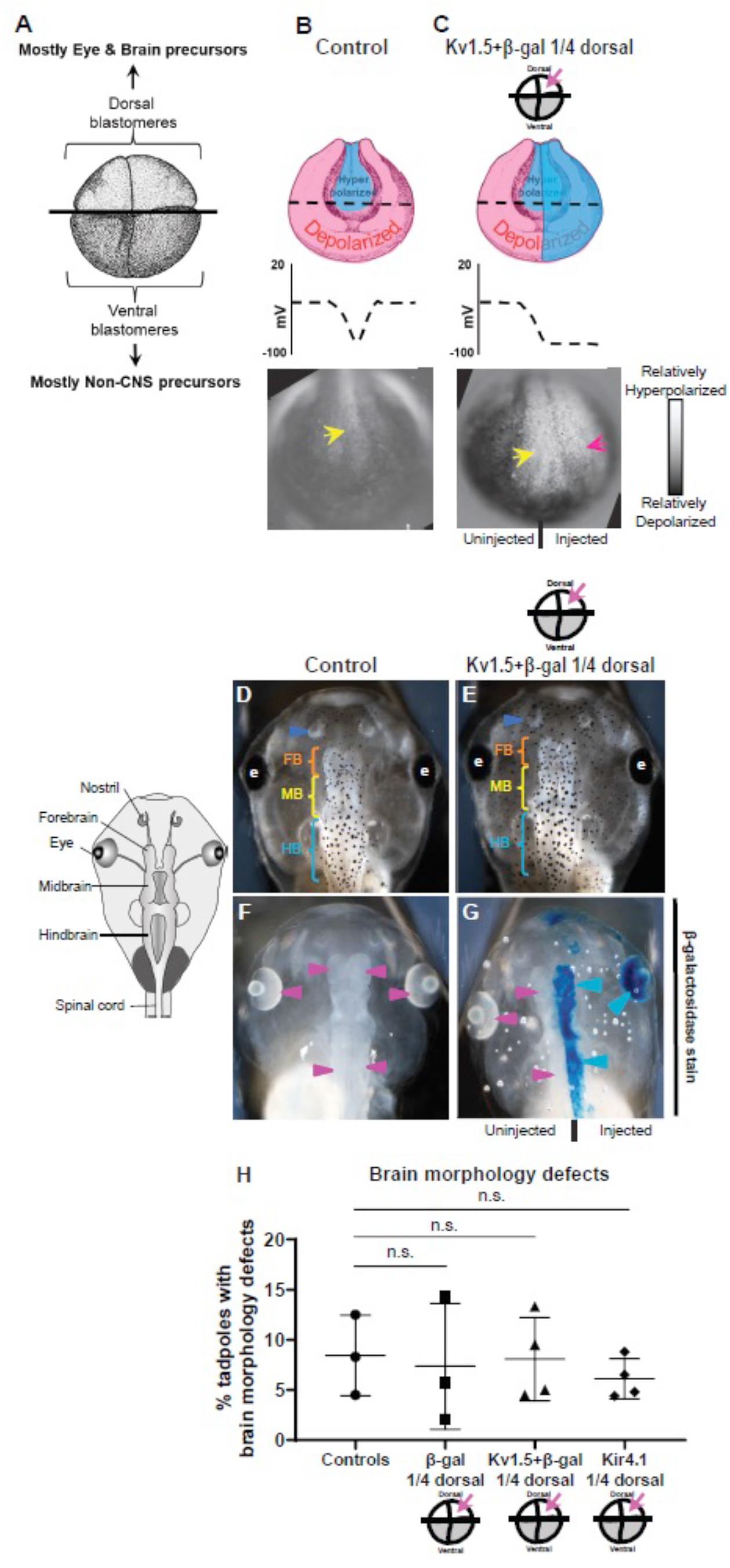
Perturbation of voltage in one half of an embryo, creating a step function voltage pattern, does not cause defects in brain morphology. (**A**) Illustration of *Xenopus* embryo at stage 3 (four-cell), indicating two dorsal blastomeres as main precursors of eye and brain and two ventral blastomeres as main precursors of non-neural tissues ^118,120^. (**B**) Normal voltage pattern in control (uninjected) stage ~15 embryo *Xenopus* embryos. Top: illustration; bottom: representative image of CC2-DMPE:DiBAC voltage reporter dyes-staining, showing characteristic hyperpolarization in the neural plate (yellow arrows) and surrounding depolarized ectoderm ^88,91,93^ (N=6). (**C**) Step function voltage pattern at stage ~15 in embryos microinjected with *Kv1.5 + β-galactosidase* mRNA in one dorsal blastomere at four-cell stage. Top: illustration; bottom: representative image of CC2-DMPE: DiBAC voltage reporter dyes-stained *embryos*, showing characteristic hyperpolarization in neural plate (yellow arrow) but hyperpolarized ectoderm only on the injected side (red arrow)(N=6). (**D-G**) Representative images of stage 45 tadpoles. (**D, E**) Tadpoles from uninjected (D) or injected (E) embryos. Blue arrowheads indicate intact nostrils, orange brackets indicate intact forebrain (FB), yellow brackets indicate intact midbrain (MB), cyan brackets indicate intact hindbrain (HB), and intact eyes (e). (**F, G**) β-galactosidase expression assessed using X-Gal (blue) in bleached tadpoles. In injected tadpoles (G), X-gal staining is evident in the eye and brain on the injected side (blue arrowheads) but not the uninjected side (magenta arrowheads), validating our targeting of mRNA microinjection. There was no X-gal staining (magenta arrowheads) in uninjected tadpoles (F) (N>10 tadpoles per experimental group). (**H**) Quantification of brain morphology defects in stage 45 tadpoles under different injection conditions demonstrates no significant differences among any of the injection conditions. Percentage of tadpoles with brain defects for each experimental group are: Controls – 8%, β-galactosidase – 7%, Kv1.5 + β-galactosidase – 8%, and Kir4.1 – 6%. Data are mean ± SD, n.s.= nonsignificant (One-way ANOVA with Tukey’s post-hoc test for n=3 independent experiments with N>50 embryos per treatment group per experiment).

These results confirm the model’s first prediction (Fig. 10A): that certain specific deviations from the normal endogenous voltage pattern responsible for brain development are well tolerated and do not result in large-scale brain morphology defects.

### 6. Empirical testing of prediction two: sharpening the voltage contrast pattern by reducing the number of neural plate hyperpolarized cells while minimally changing the overall pattern resulted in brain morphology defects

To test the model’s second prediction (Fig. 10B), we used the same strategy, but in this case injecting a well-characterized dominant-negative K_ATP_ construct (DNKir6.1p), which is known to inhibit endogenous K_ATP_ channels and cause depolarization ^20,31,121^. In comparison to control embryos, which exhibit the characteristic voltage contrast pattern and normal brain morphology^81^ (Fig. 12D &H), *DN-K_ATP_ + β-galactosidase* mRNA microinjection depolarized only the injected half of the neural plate and surrounding ectoderm (Fig. 12B-C), decreasing the number of hyperpolarized cells in the neural plate to create a narrower central band of hyperpolarized cells while maintaining the overall voltage contrast pattern (Fig. 12B-C). Our intuition (again based on our previous manipulations of the voltage pattern *in vivo*) was that this compressed, but qualitatively similar, pattern would still be sufficient to direct normal brain morphology. Once again, our results defied expectation and instead confirmed the model’s prediction. At stage 45, *DN-K_ATP_ + β-galactosidase* mRNA-injected tadpoles had significant brain morphology defects, but only on the injected side (Fig. 12E & H). β-galactosidase staining of stage 45 tadpoles confirmed intended tissue targeting (Fig. 12F-H).

**Figure 12:**
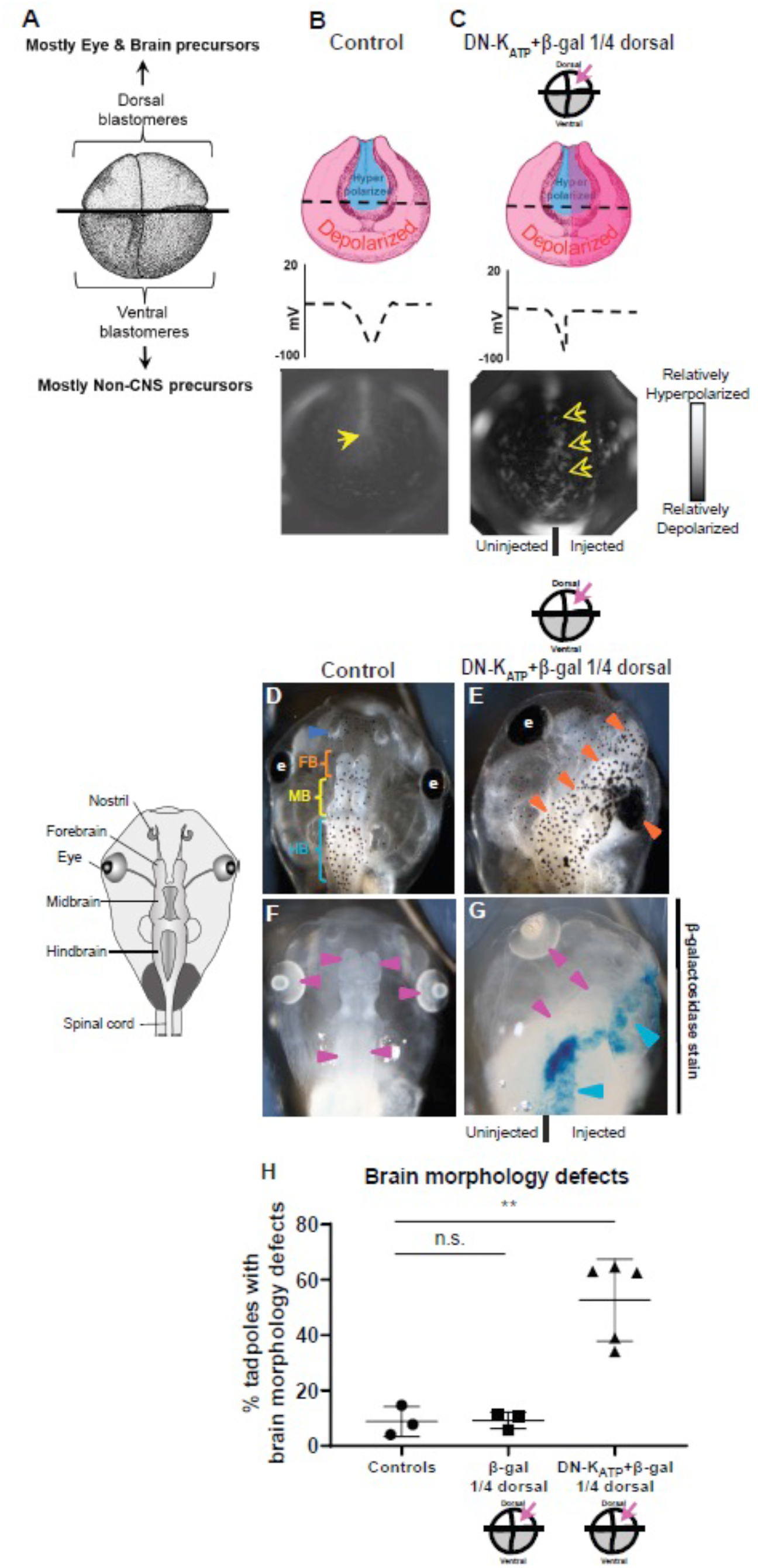
Perturbation of voltage to decrease neural plate hyperpolarized cells while minimally changing the overall voltage pattern leads to significant brain morphology defects. (**A**) Illustration of *Xenopus* embryo at stage 3 (four-cell) indicating two dorsal blastomeres as main precursors of eye and brain and two ventral blastomeres as main precursors of non-neural tissues ^118,120^. (**B**) Endogenous voltage pattern in control (uninjected) stage ~ 15 embryos. Top: illustration; bottom: representative image of embryo stained with CC2-DMPE:DiBAC voltage reporter dyes, showing characteristic hyperpolarization in the neural plate (yellow arrows) and surrounding depolarized ectoderm ^88,91,93^ (N=6). (**C**) Altered voltage pattern in stage ~ 15 embryos microinjected with *DN-K_ATP_ + β-galactosidase* mRNA in one dorsal blastomere at four-cell stage. Top: illustration; bottom: representative image of injected embryo stained with CC2-DMPE: DiBAC membrane voltage reporter dyes showing reduced region of hyperpolarization in neural plate on the injected side (empty yellow arrow). (N>6 embryos per experimental group). (**D-G**) Representative images of stage 45 tadpoles. (**D, E**) Tadpoles from uninjected (D) or microinjected with *DN-K_ATP_ + β-galactosidase* mRNA (E) embryos. Blue arrowheads indicate intact nostrils; orange, yellow, and cyan brackets indicate intact forebrain (FB), midbrain (MB), and hindbrain (HB), respectively. (e) indicates intact eyes, and orange arrowheads indicate mispatterned brain and eye. (**F, G**) β-galactosidase expression assessed using X-Gal (blue) in bleached tadpoles either left uninjected (controls) or co-injected with *DN-K_ATP_ + β-galactosidase* mRNA. There was no X-gal staining (magenta arrowheads) in uninjected tadpoles (F). In injected tadpoles (G), β-galactosidase was observed in the eye and brain on the injected side (blue arrowheads) but not the uninjected side (magenta arrowheads), confirming expected targeting of microinjected mRNAs. (N>10 tadpoles per experimental group). (**H**) DN-K_ATP_ + β-galactosidase injection significantly increased the percentage of stage 45 tadpoles with brain morphology defects: uninjected or β-galactosidase controls – both 9%; DN-K_ATP_ + β-galactosidase – 53%. Data are mean ± SD, **p<0.01, n.s.=non-significant (One-way ANOVA with Tukey’s post-hoc test for n=3 independent experiments with N>50 embryos per treatment group per experiment).

These results confirm the model’s second prediction (Fig. 10B): that, in addition to the shape of the voltage pattern, the proportion of depolarized and hyperpolarized cells that form the pattern is also a crucial factor in regulating embryonic brain development.

## Discussion

A central goal of developmental biology is to understand how multiple modalities (e.g., biophysical and transcriptional) interact to reliably produce the correct target morphology of complex organs. Bioelectric signaling is becoming increasingly recognized as an important component of developmental control mechanisms ^17–19^, especially as concerns craniofacial patterning ^28,29^. While great strides have been made in dissecting cellular-level transduction mechanisms and specific transcripts downstream of voltage-mediated signaling, much remains to be learned about the bioelectric code ^80^: the mapping of tissue-scale bioelectric prepatterns to specific organs, and the interpretation of bioelectric gradients by cellular collectives ^55,57,58,78,93,122^.

Specifically, a key question concerns how downstream morphogenetic decisions (such as formation of eyes ^20^, patterning of the anterior-posterior ^59,123^ and left-right ^30,117^ axes, and size control ^37–40^) are triggered by voltage patterns across cell fields (not simply the cell-autonomous control of transcription by the cell’s own V_mem_). In other words, diverse aspects of organ size and shape, driven by cell behaviors (migration, differentiation, apoptosis, and gene expression), are triggered by *specific multiscale patterns* of membrane potential. How can tissue read a large-scale pattern as input and determine whether it is correct or not?

This is the same problem faced by nervous systems: behavioral modules must be triggered by complex patterns of inputs in the retina, not the states of individual retinal cells. The tissue bioelectric system is an ancient version of a complex set of circuits which began by controlling the traversal of body configuration through anatomical morphospace ^3^ and eventually evolved to solve the same problem in behavioral spaces ^124^. Given that tissue bioelectric signaling is thought to be the evolutionary origin of the brain and its multiscale, top-down pattern recognition capacities ^1,9^, it is perhaps not surprising that development features some of the same computational tasks ^125^.

We sought to develop a specific model of the interplay between ionic control mechanisms and gene-regulatory networks that would identify, in a constructivist manner, dynamics sufficient for native gene-regulatory circuits to reach the correct state downstream of recognizing the appropriate organ prepattern. The *Xenopus* brain is an ideal context for such work because both the correct pattern of bioelectric signals for normal brain behavior ^88,89,91–93^ and the transcriptional cascades required for brain development ^97–99^ are known.

The minimal recurrent dynamical model described here, designed using supervised machine learning techniques, recapitulates the experimentally observed phenomenon of bioelectric control of morphogenesis in the nascent *Xenopus* embryo: only certain appropriately shaped voltage contrast patterns across the neuroectoderm tissue lead to the correct development of the brain. Analysis of the information-processing mechanisms underlying the model’s pattern discrimination behavior revealed that it employs a feedforward-like control mechanism (Fig. 7) where the spatial voltage pattern influences a relatively more dominant control on gene expression, rather than the other way around, despite the complex feedback connectivity between the bioelectric patterns and the genes (Fig. 3). Further analysis revealed a unique oscillatory mechanism with which different regions of the tissue influence gene expressions in distant cells at different points in time (Fig. 8). The statistical properties of this mechanism predicted that other spatial voltage patterns with atypical ratios of hyperpolarized versus depolarized cells in the neural plate tissue could also result in a normal brain (Fig. 10) – predictions that were experimentally verified (Figs. 11,12).

Our model was designed to be minimal and phenomenological, hence it cannot offer detailed insights into the microscopic mechanisms of the cells that may also be contributing to processing the bioelectric patterns. Even though this may be considered as a limitation of the model, it may also be considered as a strength in that it helps focus on the factors that may be relevant to the problem at hand (bioelectric pattern recognition in our case). The other limitation is that it may not be easily amenable to incorporating additional biological knowledge; this can only be accomplished by training the model from scratch. Lastly, our analysis offers only one possible set of mechanisms that the neural ectoderm might be employing, though experimental validation of their predictions provides compelling support. It may be interesting in the future to map the space of all possible mechanisms in a way that could even throw light on the evolutionary aspects of the system.

Our model demonstrates how bioelectric control of gene expression can extend beyond the scope of a single cell ^49^ by showing that the macroscale shape of bioelectric patterns across a tissue can direct gene expressions in the (microscale) cells comprising it. Importantly, the model was not micro-managed to implement all of the features we found; our analysis of this model revealed numerous surprising emergent features, thus being also an example of digital embryogeny ^126,127^. Our analysis revealed interpretable spatiotemporal information-integration mechanisms that the model employs to both recognize and discriminate between the various voltage patterns, as depicted in Figs. 6–9.

Some of the unique features of the mechanism include functional specialization and long-distance influence. There are three different ways in which functional specialization manifests in our model. First, certain cell positions are more important in interpreting bioelectric patterns (Fig. 6). This suggests that collective cell decision making in this context occurs via a partially distributed system, which allows the system to be robust to external perturbations and at the same time be highly plastic, since change in a few key cell positions can drastically change the interpretation of patterns. This is supported by our *in vivo* experiments that demonstrate that while drastic changes in the bioelectric pattern still result in normal brain morphology (Fig. 12) relative minor changes could significantly alter the outcomes (Fig. 13).

**Figure 13.**
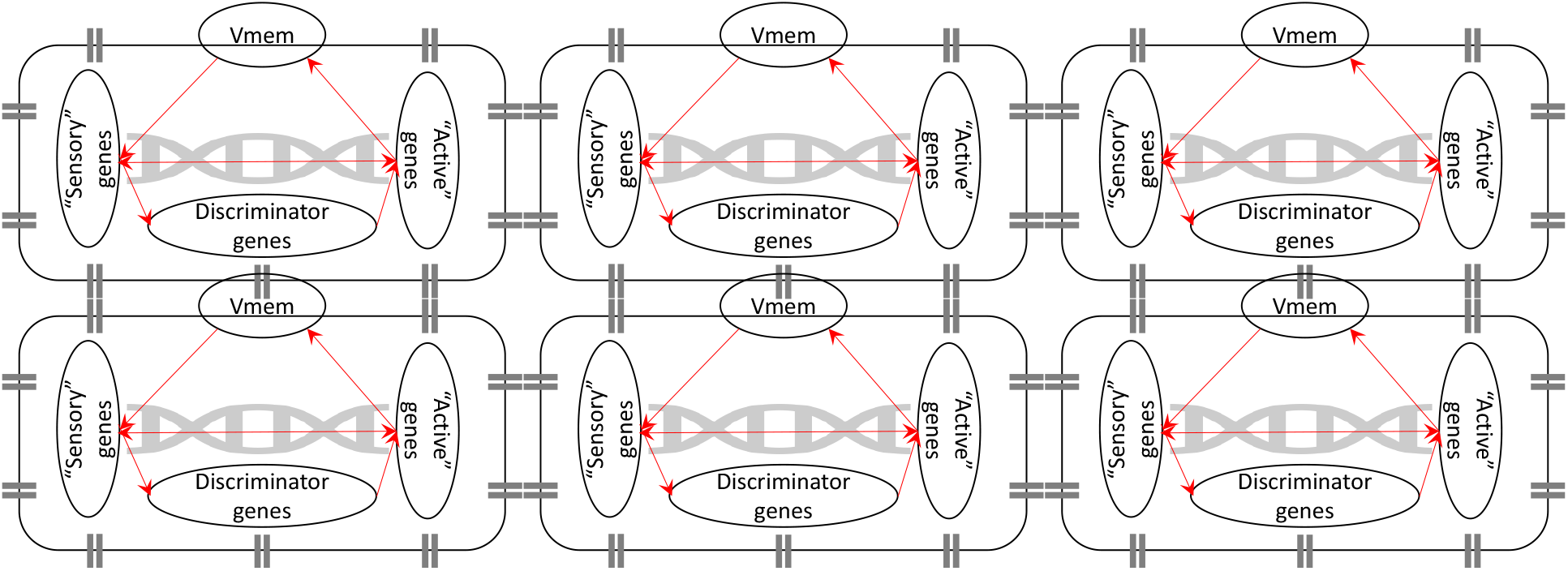
A schematic illustration of an active inference perspective of the voltage pattern discrimination. Each block represents a cell containing ion channels and gap junctions on the periphery and a gene regulation network at the core. The neural plate-ectoderm circuit is hypothesized as analogous to the active inference circuit thought to underlie cognition in neural systems^149^. Here, voltage pattern is conceptualized as the external (hidden) variable that the discriminator gene internally represents and infers via the Markov blanket consisting of the “sensory” and the “active” genes. The graph depicted inside each cell need not match the structural graph of the model (Fig 4); it could also dynamically emerge by virtue of nonlinearity. The inference mechanism existing inside a single cell could in principle scale up to the level of the tissue that would then be said to be performing collective active inference ^154^ (not illustrated). Such a collective active inference mechanism could lie at the heart of the neuroectoderm voltage pattern discrimination behavior.

Moreover, different genes are sensitive to different spatial scales of the bioelectric pattern, ranging from the level of the tissue to that of the single cell (Fig. 9). While genes are conventionally viewed as coding for scales of the organism ranging from proteins to phenotypic features and diseases, our model flips this perspective and highlights the scales to which they could be *receptive*, at least within the confines of a multicellular tissue. This has implications for gene therapy. For instance, it may be possible to target certain genes not directly but indirectly through larger scale features like bioelectric potentials that may be relatively easier to manipulate (more discussion along these lines below).

Finally, the proportion of depolarized to hyperpolarized cells in the tissue may determine how the voltage pattern gets processed in terms of gene expression (Figs. 10, S5). That is, the extent to which the voltage of a cell influences the gene expression in other cells depends on the type of polarization of other cells. This is validated by our *in vivo* experiment where a slight change in proportion of polarized cells while keeping the overall bioelectric pattern same results in major brain development deformities. Particularly of interest is the observation that the depolarized (non-neural ectoderm) cells are more important in this collective cell decision making process than the hyperpolarized (neural ectoderm) cells. This means that developmental patterning of any tissue or organ is as much dependent on cells in the surrounding embryo as on the cells making up that tissue and organ. This in hindsight makes sense as no embryonic tissue develops in a vacuum but instead is part of a whole organism and all tissue developments have to be somewhat coordinated for the proper development of the whole organism.

Another central feature of the information-processing mechanism that our model employs is long-distance influence (Fig. 8), showing how genes can be responsive to long-distance influence from far away cells ^128^. Not only is the responsiveness of genes not static but it is also dynamically fluctuating between local responsiveness and distant responsiveness. This adds another layer of complexity to gene expression and regulation especially during active morphogenesis, where gene expression at any given time can no longer be considered as a static readout of the state of that cell but should be viewed as a culmination of several influences, local and distant, perhaps similar to that of input integration on a neuron.

Yet another unique feature of our model is its simplified control mechanism (Fig. 7), where the voltage pattern behaves as if it directly controls the discriminator gene, in effect masking the recurrent connectivity amongst the genes (a simple analog of this setting is described in SI. 9). This feature is analogous to the concept of “canalization” that’s known to often endow biological systems with the ability to shield itself from random mutations ^129^. In the context of complex dynamical systems it translates into a phenomenon where only a subset of the nodes or pathways in the network actually direct the system to its final attractor state for a given initial state ^130,131^. Our model also exhibits a similarly canalizing relationship between the voltage pattern and the discriminator gene expression in that the former is sufficient to determine the asymptotic expression of the latter. Moreover, we’ve described this relationship in formal terms and characterized it as existing at the second-order level (Fig. 7), as indicated by a dominant Hessian tensor. This mathematical understanding of canalization in the specific context of the relationship between bioelectricity and gene expression is crucial to advance our understanding of the bioelectric control of morphogenesis in *X*enopus and possibly other biological systems.

Our model generated empirically testable predictions (Figs. 10–13) that were surprising. For instance, altering the bilaterally symmetrical endogenous bioelectric pattern, which is known to be required for normal brain development ^91,93^, by disrupting half of it still led to normal brain development *on both sides*. This confirms the ability of the combined bioelectric-GRN system to collectively achieve outcomes despite perturbations by leveraging correct information on one side to accommodate informational defects on the other side. This especially makes sense in light of the observation that the depolarized cells in our model are more influential than the hyperpolarized cells (Fig. S5). On the other hand, changes in the proportion of polarized cells, while maintaining the overall bioelectric pattern, led to major brain development defects. This is also counter-intuitive because the overall bioelectric pattern is maintained and so it should have led to normal brain patterning. These observations moreover indicate the robustness of this system to perturbations: as long as critical information integration and interpretation nodes are maintained the correct target morphology will be achieved. These results suggest that a useful perspective is to adopt the lens of how the cells themselves “see” and interpret surrounding signal patterns. What might seem an obvious pattern to an observer scientist might not correctly reflect the underlying salience to the cell group, given the computations and interpretations being made by the collective. This underscores the importance of making models like this and analyzing them for effective triggers of change (and input invariants), which may not be otherwise obvious.

Finally, we can ask if the physical boundaries of a given cell in our multicellular model (Fig. 1F) are relevant and to what extent. Are there larger or smaller *effective* boundaries cutting across different clusters of cells and genes that are more meaningful for the pattern discrimination problem? It has been hypothesized, in the context of cancer and morphogenesis, that when active biological units such as cells come together in informationally-connected groups ^132–134^, the computational boundary demarcating a coherent “individual” could scale up from single units to the collective. Large-scale phenomena such as morphogenesis have been described as behaviors of a collective intelligence in anatomical morphospace ^3,135,136^. Could our neuroectoderm tissue similarly possess functional boundaries larger than that of a single cell in ways that could facilitate pattern discrimination at the tissue level? Our analysis has already provided hints to support this view, where small clusters of cells seem to act as coherent *modules* in terms of the constituent cells’ similarity in behavior during the spatiotemporal integration process (Fig. 8). Moreover, more than one such instructive causal nexus exists in the tissue – these may be separated in physical space but interact with each other at certain times (Fig. 8) – an indication that they may be closer in “physiological space” even if distant in physical space.

The existence of two different modalities in our model, namely bioelectricity and genetics, with a simplified feedforward-like relationship between them raises interesting possibilities concerning the control of the latter by the former. In particular, our model demonstrates that behavior in the higher-dimensional genetic space could be controlled by a lower-dimensional bioelectric space; the former is larger than the latter by a factor of *n_g_*, the number of genes. This capacity for efficient control of gene expression is related to canalization discussed above. The traditional sense of the term implies that it’s sufficient to specify the state of a subset of the nodes to guarantee the convergence of the state of the entire dynamical system to a specific attractor ^130,137^. In our bioelectric/genetic hybrid model, we posit that it is the bioelectric state that effectively acts as the independent external input to the genetic variables, thus acting as a control knob for the gene expression pattern. Of course, the alternative is to directly control the genetic state simply by freezing it at a desired state or specifying some other state that can guarantee convergence to the desired state – such an approach would be less efficient than bioelectric control due to the fewer variables involved. Analysis revealed that, in our model, the voltage pattern does indeed control gene expression in an apparently feed-forward manner despite the presence of recurrent connectivity between them (Fig. 10). Indeed, biological systems employ such mechanisms for morphological control; for example, voltage prepatterns can be modulated to alter the number of heads in planaria ^36^ or the craniofacial features of frog embryos^28^. By virtue of the relative simplicity of control, this allows biological systems, and perhaps also evolution, to encode target morphologies in a potentially more controllable software-like system (such as bioelectricity) compared to the more complex microscopic “hardware” (genetically-encoded elements) which features difficult inverse problems ^138^.

Our findings may also have implications for the emerging field of basal cognition ^139^. Here, we adopt the perspective of the “cognitive lens” ^3,139–141^ – a prescription for a unified view of biological systems as integrated wholes that may leverage universal computational principles, regardless of their biochemical compositions or scale. The competencies identified in our model can be characterized as instances of very primitive cognition, when viewed as generic sensing/actuation and other information-processing loops that biological organisms, whether single or multicellular, neural or non-neural, employ to navigate diverse problem spaces for the purpose of survival, growth and reproduction ^140,142,143^. We leveraged the causal integration analysis framework ^103^ to decipher such higher level characterizations from the underlying mechanistic details and recast them in terms of basal cognition. For example, the neural plate genes could be viewed as paying “attention” to the voltage pattern while ignoring the inputs from other genes (Fig. 7) and “attention-shifting” (Figs. 8,9) to integrate information from various regions and spatial scales of the tissue in order to “perceive” (Fig. 6) the endogenous voltage pattern by recognizing its characteristic features. Here, we adopt the definitions of attention and perception from Ref ^143^, where attention is the “ability to selectively attend to a state of affairs to the exclusion of the others” and perception is the “capacity to sense and recognize (re-cognize) existentially salient features of the external and internal milieux”. This high-level perspective of our model offers a way to decompose microscopic biochemical behavior into “cognitive modules”, which suggest that future bioengineers and workers in regenerative medicine could deploy tools of behavior science and cognitive science to exert optimal control over such systems in biomedical contexts ^9,144–146^. In other words, these perspectives are useful not only in understanding and generating hypotheses about the “software” that the modeled biological system might employ but also for functionally reprogramming it to achieve alternative outcomes, especially when the model generates correct predictions. For example, we may be able to trick the tissue into misperceiving the endogenous pattern by interfering with the attention mechanism – a form of minimal illusion. The precise methods for accomplishing this would be the subject matter of future research but are already underway given the discoveries of memory and learning in gene-regulatory networks ^146,147^.

Another theoretical framework that’s becoming increasingly popular in characterizing cognition-like characteristics of a wide variety of biological systems, neural or otherwise, is active inference^148^. According to this framework, any random dynamical system that contains an “inside” and an “outside” separated by a “Markov blanket”, consisting of “sensory” and “active” states, could be shown to be performing Bayesian inference and hence is cognitive ^149^. That is, the internal states of such a system could be shown to be predicting the external causes of the sensory states and at the same time actively causing the external states through action. This formalism has already begun to be used to understand developmental patterning ^150–153^. Our model is a first step towards a research program that quantitatively casts the embryonic neural plate-ectoderm circuit as an active inference system, where the discriminator genes infer the voltage pattern of the tissue via a layer of “blanket” genes (Fig. 13).

Taken together, these results weave together the components of an exciting emerging field, including developmental biophysics, dynamical systems, and machine learning. Future advances in computational perspectives on the activity of multi-scale biological systems will reveal both mechanistic and cognition-based perspectives that optimize insight, prediction, and control. These frameworks will be an essential component of the future roadmap to exploit the innate competencies of biological systems as information-processing agents, thus overcoming the many fundamental barriers facing purely bottom-up molecular approaches. Complementary fusion of both approaches will greatly potentiate the discovery of interventions for regenerative medicine and synthetic bioengineering.

## Acknowledgements

We thank Erin Switzer and Rakela Colon for *Xenopus* husbandry and general laboratory assistance. We thank Joan Lemire for assistance with cloning of cDNAs into frog expression vectors. We thank many members of the Levin lab for helpful discussions, as well as Susan Lewis and Julia Poirier for assistance with the manuscript.

## Author contributions

S.M. performed computational model development and analysis, V.P.P. performed in vivo experiments, S.M., V.P.P, and M.L. designed experiments, interpreted data and wrote the manuscript.

## Supplementary Information

### 1. Parameter details

**Table S1.**
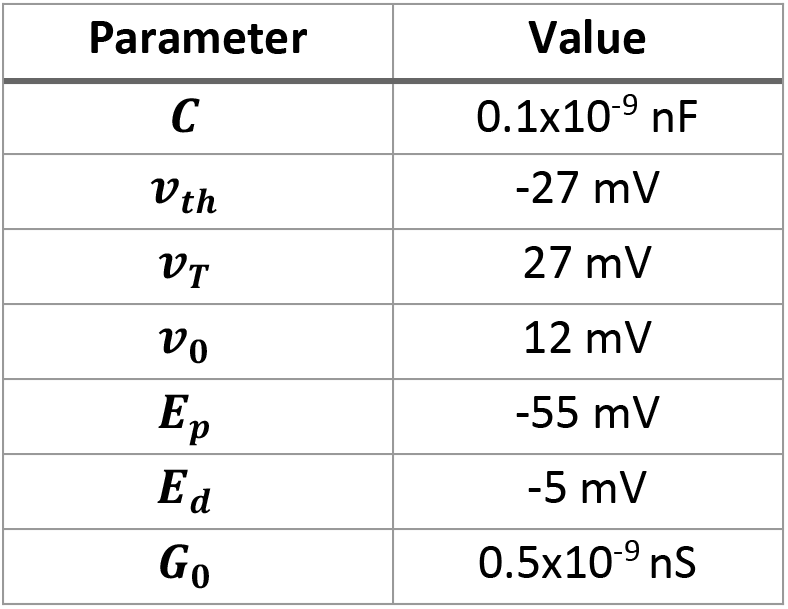
Bioelectric constants

**Table S2.** ***W**^g→g^* values (Arbitrary Units)

| | $g_{i,1}$ | $g_{i,2}$ | $g_{i,3}$ | $g_{i,4}$ | $g_{i,5}$ | $g_{i,6}$ |
| --- | --- | --- | --- | --- | --- | --- |
| $g_{i,1}$ | 0 | 12.1406 | 24.7507 | 0 | -21.6979 | 0 |
| $g_{i,2}$ | 5.867 | 0 | 19.0281 | 8.2776 | 2.7572 | -15.7728 |
| $g_{i,3}$ | -5.2364 | 16.79 | 0 | 40.9563 | -276.7888 | 8.2963 |
| $g_{i,4}$ | 9.4655 | -12.4545 | 28.1367 | 0 | 5.2432 | 11.7852 |
| $g_{i,5}$ | -13.343 | 12.2471 | 20.0379 | -4.9928 | 0 | 0 |
| $g_{i,6}$ | -15.7708 | -8.6491 | 16.8677 | 2.9967 | -18.7133 | 0 |

**Table S3.** ***W**^g_N_→g^* values; (*i,j*) *are neighboring cells* (Arbitrary Units)

| | $g_{j,1}$ | $g_{j,2}$ | $g_{j,3}$ | $g_{j,4}$ | $g_{j,5}$ | $g_{j,6}$ |
| --- | --- | --- | --- | --- | --- | --- |
| $g_{i,1}$ | -0.9017 | 5.2314 | 0 | 0 | -56.0558 | 9.4501 |
| $g_{i,2}$ | 0 | 10.9099 | 34.0221 | 5.5974 | 8.4497 | 5.6936 |
| $g_{i,3}$ | 13.4256 | -4.4949 | 26.9793 | 29.3938 | -194.6024 | 13.8529 |
| $g_{i,4}$ | 4.8829 | 0 | 0 | 23.1135 | 2.5301 | -4.1686 |
| $g_{i,5}$ | 8.2243 | -1.5515 | 1.8714 | -0.6805 | -3.3403 | -11.0029 |
| $g_{i,6}$ | 0 | -5.8101 | 0 | 15.6676 | -1.8589 | 13.0818 |

**Table S4.** ***W**^v→g^* values (Arbitrary Units)

| | $g_{i,1}$ | $g_{i,2}$ | $g_{i,3}$ | $g_{i,4}$ | $g_{i,5}$ | $g_{i,6}$ |
| --- | --- | --- | --- | --- | --- | --- |
| $v_i$ | 27.7108 | 17.2487 | 6.8623 | -28.9754 | 35.5804 | 25.6036 |

**Table S5.** ***W**^g→v^* values (Arbitrary Units)

| | $g_{i,1}$ | $g_{i,2}$ | $g_{i,3}$ | $g_{i,4}$ | $g_{i,5}$ | $g_{i,6}$ |
| --- | --- | --- | --- | --- | --- | --- |
| $v_i$ | 0 | 3.2776 | 0 | 0 | 1.9991 | 0 |

**Table S6.** ***b_g_*** values (Arbitrary Units)

| $g_{i,1}$ | $g_{i,2}$ | $g_{i,3}$ | $g_{i,4}$ | $g_{i,5}$ | $g_{i,6}$ |
| --- | --- | --- | --- | --- | --- |
| -11.3735 | -3.6548 | -14.2749 | -8.385 | -1.6196 | -6.2071 |

**Table S7.** ***τ_g_*** values (Arbitrary Units)

| $g_{i,1}$ | $g_{i,2}$ | $g_{i,3}$ | $g_{i,4}$ | $g_{i,5}$ | $g_{i,6}$ |
| --- | --- | --- | --- | --- | --- |
| 1.04e-01 | 1.66e+01 | 2.47e+01 | 5.66e+00 | 1.89e+01 | 1.20e-02 |

**Table S8.** Other scalar parameter values (Arbitrary Units)

|  |  |
| --- | --- |
| $\tau_d$ | 1.3349 |
| $\tau_r$ | 1.2819 |
| $\tau_v$ | 0.5071 |
| $b_v$ | 0.0 |
| $u$ | 4.7180 |

### 2. Details of the machine-learning algorithms

For the GD, we employed “resilient backpropagation” that relies only on the sign (not the magnitude) of the gradients for updating the parameters at every iteration, with a learning rate of 0.01.

For the microbial GA, we used a population of 40 individual “genotypes”. Each genome codes for the parameters and the meta parameters of a single instance of the model. The parameters are the literals listed in red in Fig. 2A coding the parameters of a model with a given size of the GRN and the iGRN, whereas the meta parameters, coding for the size of the GRN and the iGRN, are the following: number of genes (*n_g_*), number of GRN edges (*n_e_*), number of iGRN edges (*n_i_*), number of GRN to Vmem edges (*n_gv_*), and the number of Vmem to GRN edges (*n_vg_*). Every gene in a genotype has a value in the range [0.01,1] that is linearly mapped to the corresponding model features according to their respective ranges, as follows: weights have a range of [-16,16], timeconstants [0.1,30], bias of the GRN nodes [-16,16], bias of the Vmem node [-100 mV,0 mV], Vmem gain [0,7], *n_g_* [6,20], *n_e_* has a range of [*n_g_* – 1, *n_g_*(*n_g_* – 1)], *n_i_* has a range of 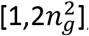, and both *n_gv_* and *n_vg_* have a range of [1, *n_g_*]. We used a “geographical selection” method where the individuals (genotypes) are placed on a 1D ring, and only geographically close individuals are picked to compete in the tournaments. The size of selection-neighborhood, known as the “deme size”, was set to 20%. The genome of the winner of a tournament was transmitted to the loser at a “crossover rate” set to 10%. The genome of the loser was then mutated at a “mutation rate” of 5% and reinserted into the population. All genotypes were randomly initialized (in the range [0.01,1]) during the first generation. This process was repeated for about 1200 generations.

Each genotype was evaluated by simulating the corresponding model and computing its performance at the end of the simulation. The initial conditions of the variables in each simulation were set as follows: *v* = −9.2mV (the unstable equilibrium point in the bistable cell) for all cells; *G_p_* = 1.0*G*_0_ for the cells in the left and the right bands and 1.8*G*_0_ for the cells in the middle band, while the constant *G_d_* was set to 1.5*G*_0_ for all the cells; the values of all other variables including *G_ij_* and *g* were set to 0. The model equations were integrated using the standard Euler method with a fixed step size of 0.01 for about 1000 steps.

### 3. Calculation of the discriminatory power of a gene

The *discrimination score*, 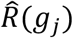, of a gene *g_j_* is defined as the change in the observed discrimination error, *R*_obs_(*g_j_*), relative to the maximum possible error, *R*_max_, where the *discrimination error* is computed as the weighted mean distance between the observed gene expression pattern and the target patterns corresponding to the endogenous (*E*), depolarized (*D*), and hyperpolarized (*H*) input Vmems, 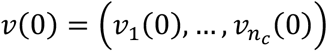:

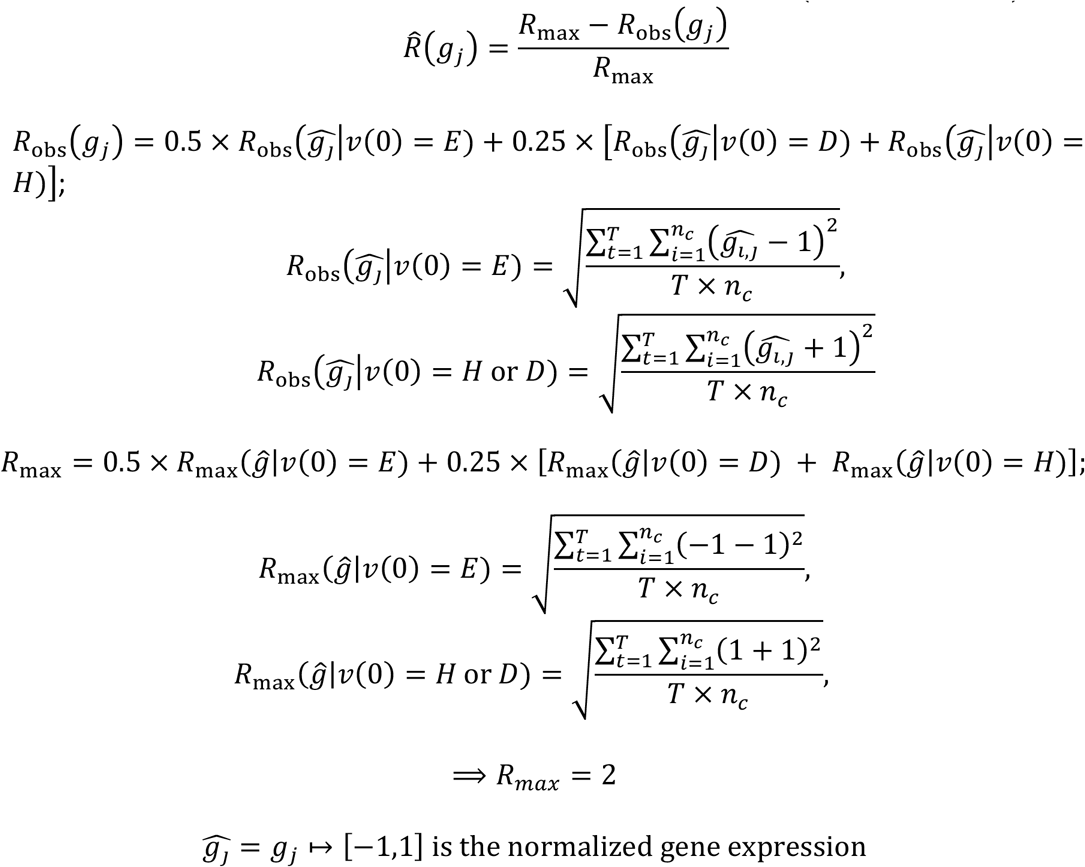

The desired target gene expression patterns are the following: all-activated (1, …, 1) for *E;* and, all-repressed (−1, …, −1) for *D* and *H*. Thus, the value of 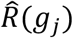 lies between 0 and 1, corresponding to the minimum and maximum possible discriminatory scores, while a value of 0.5 indicating a randomly expected score (no discrimination).

### 4. Calculation of the performance of a model

The *performance score, P*, of the model is defined as the complement of the ratio between the observed mean discrimination error, averaged over all genes, and the maximum possible discrimination error:

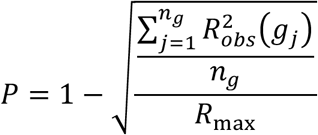

Thus, the value of *P* lies between 0 and 1, corresponding to the minimum and maximum possible performance, while a value of 0.5 indicating the expected performance of a randomly parametrized model.

### 5. Training improves the performance of an ensemble of models

**Figure S2.**
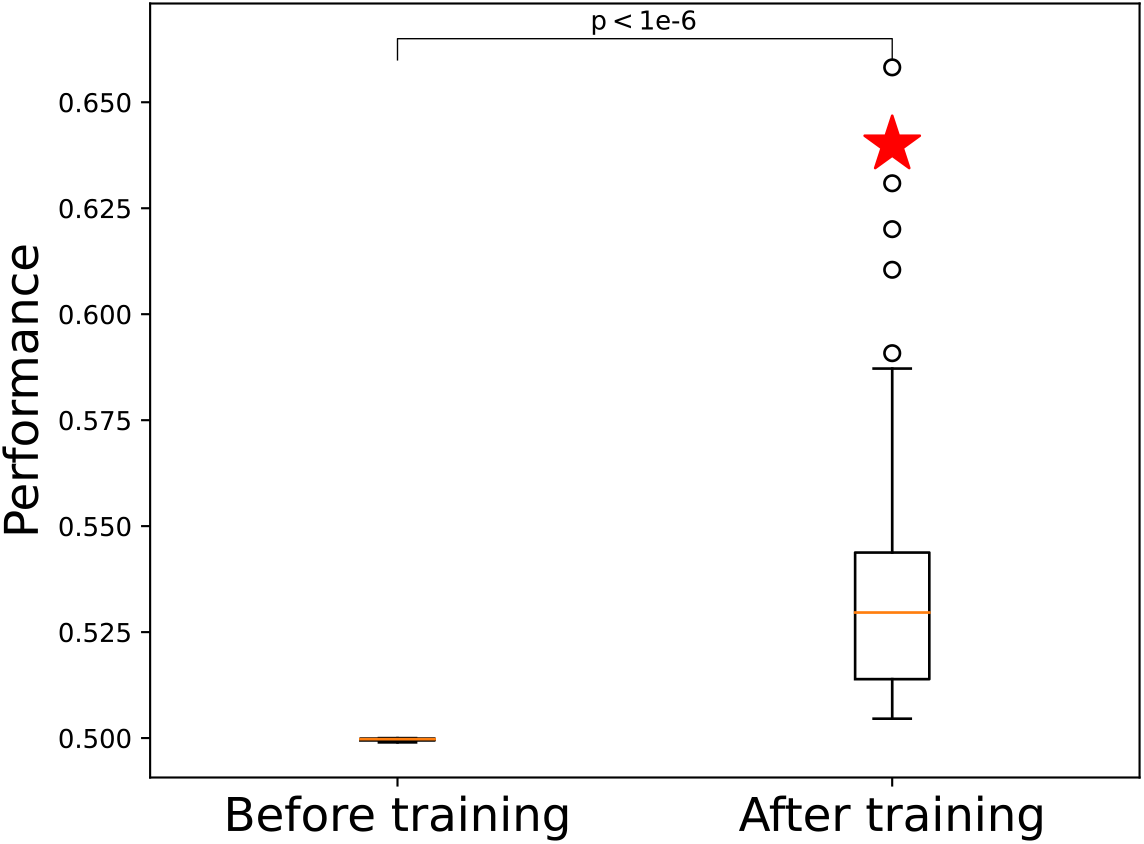
Comparison of the performances of an ensemble of models before and after training. The after-training group is significantly better than the before-training group as per t-test (p<1e-6). The red star indicates the performance of the model analyzed in the main text.

### 6. Model qualitatively recapitulates the bioelectric pattern discrimination phenomenon observed in *Xenopus* embryos

**Figure S3.**
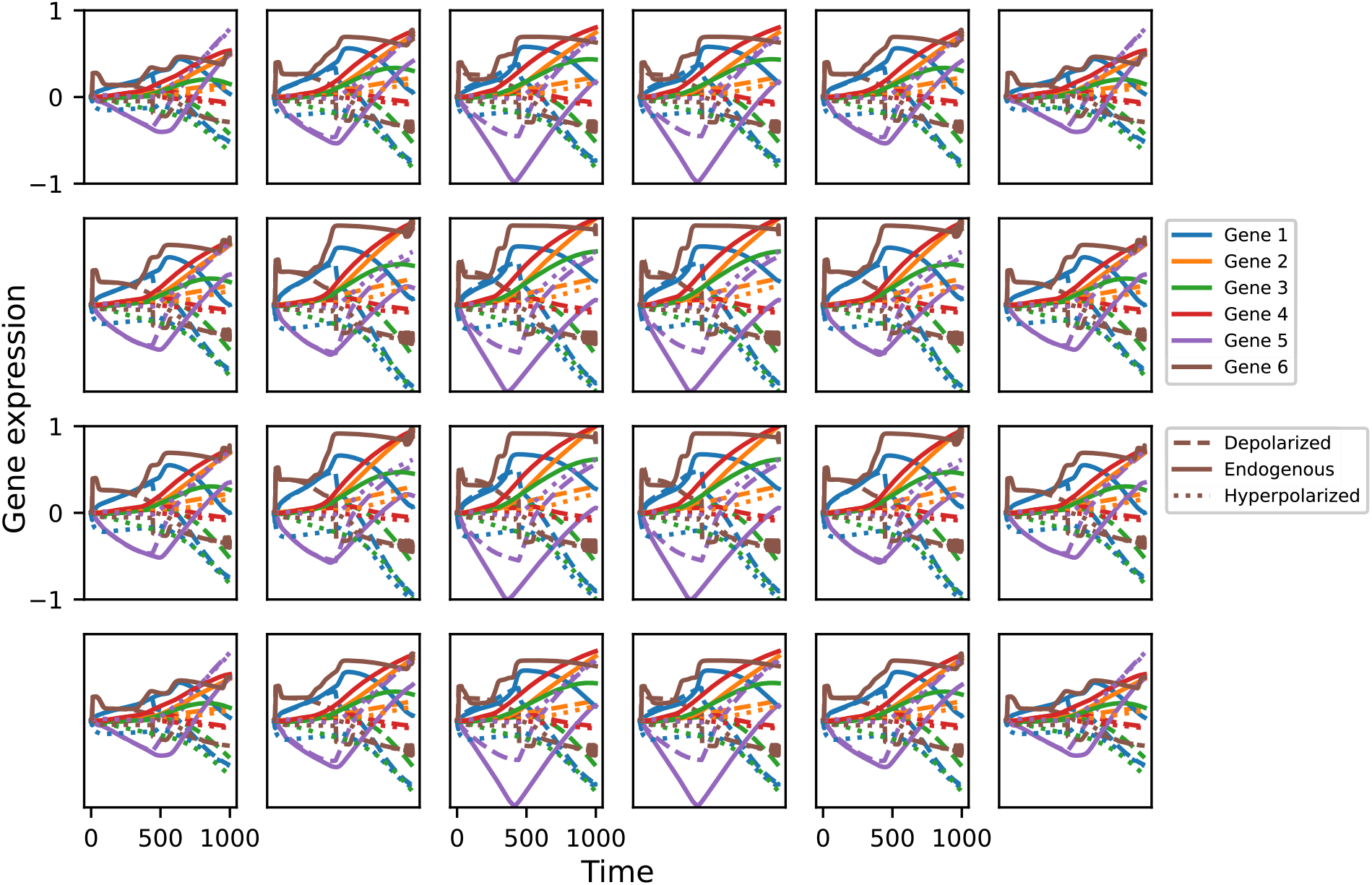
Cell-wise behavior of the individual genes while solving the pattern discrimination problem in a 4×6 tissue. Most genes, regardless of the cell, tend to activate (positive values) for the endogenous bioelectric pattern while tending to deactivate (negative values) for the aberrant patterns. It can also be noticed that while the quantitative gene expression behavior depends on the cell’s spatial location (up to bilateral symmetry across both the vertical and horizontal axes passing through the center of the tissue), there are no significant qualitative differences.

### 7. Model solves the pattern discrimination problem even in larger tissues despite not having been selected for such capability: scale robustness

**Figure S4.**
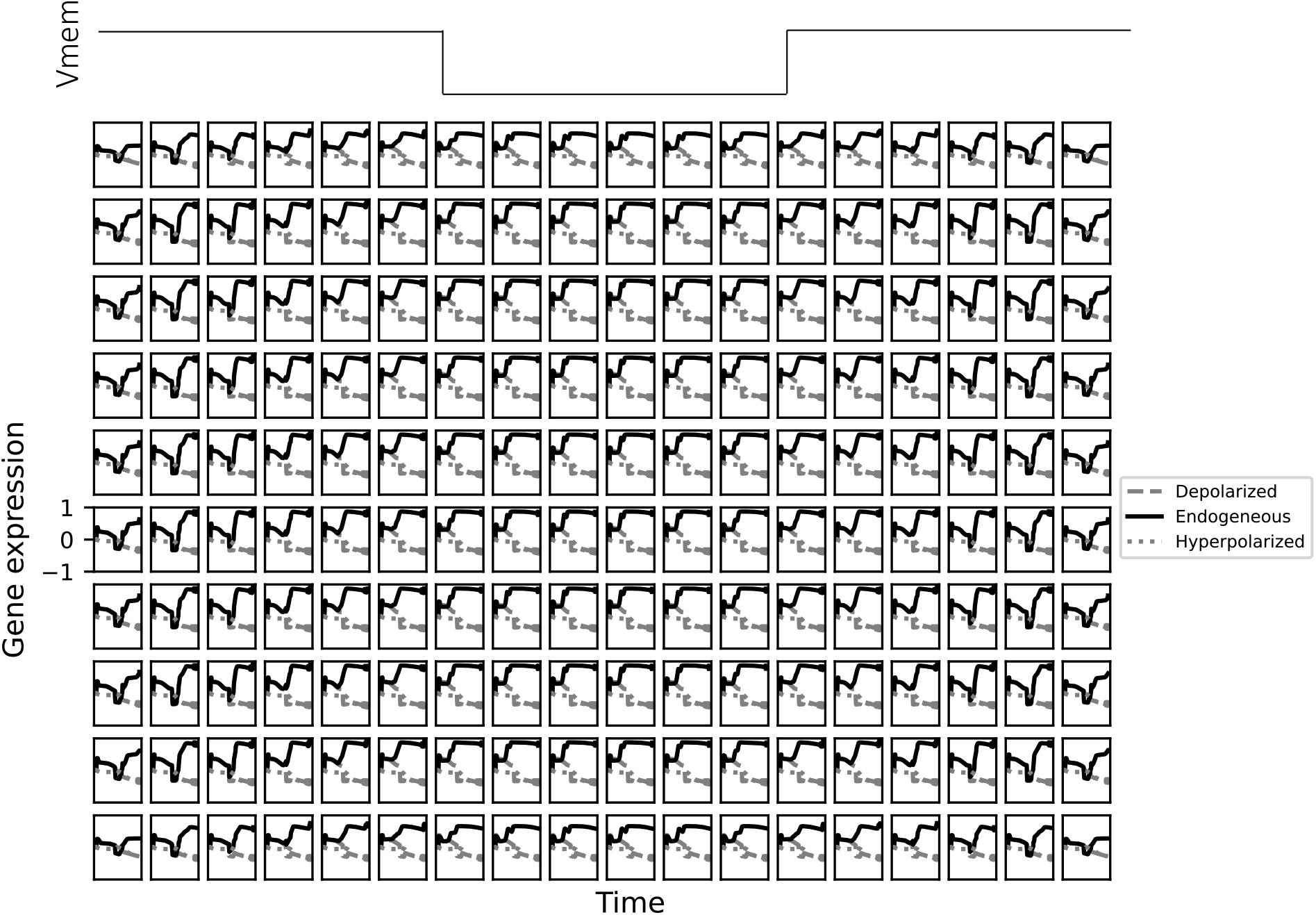
Cell-wise behavior of the discriminator gene while solving the pattern discrimination problem in a 10×18 tissue. Regardless of the cell, the discriminator gene tends to activate (positive values) for the endogenous bioelectric pattern (top) while tending to deactivate (negative values) for the aberrant patterns. It can also be noticed that while the quantitative gene expression behavior depends on the cell’s spatial location (up to bilateral symmetry across both the vertical and horizontal axes passing through the center of the tissue), there are no significant qualitative differences.

### 8. Model predicts a class of aberrant spatial bioelectrical input patterns

**Figure S5.**
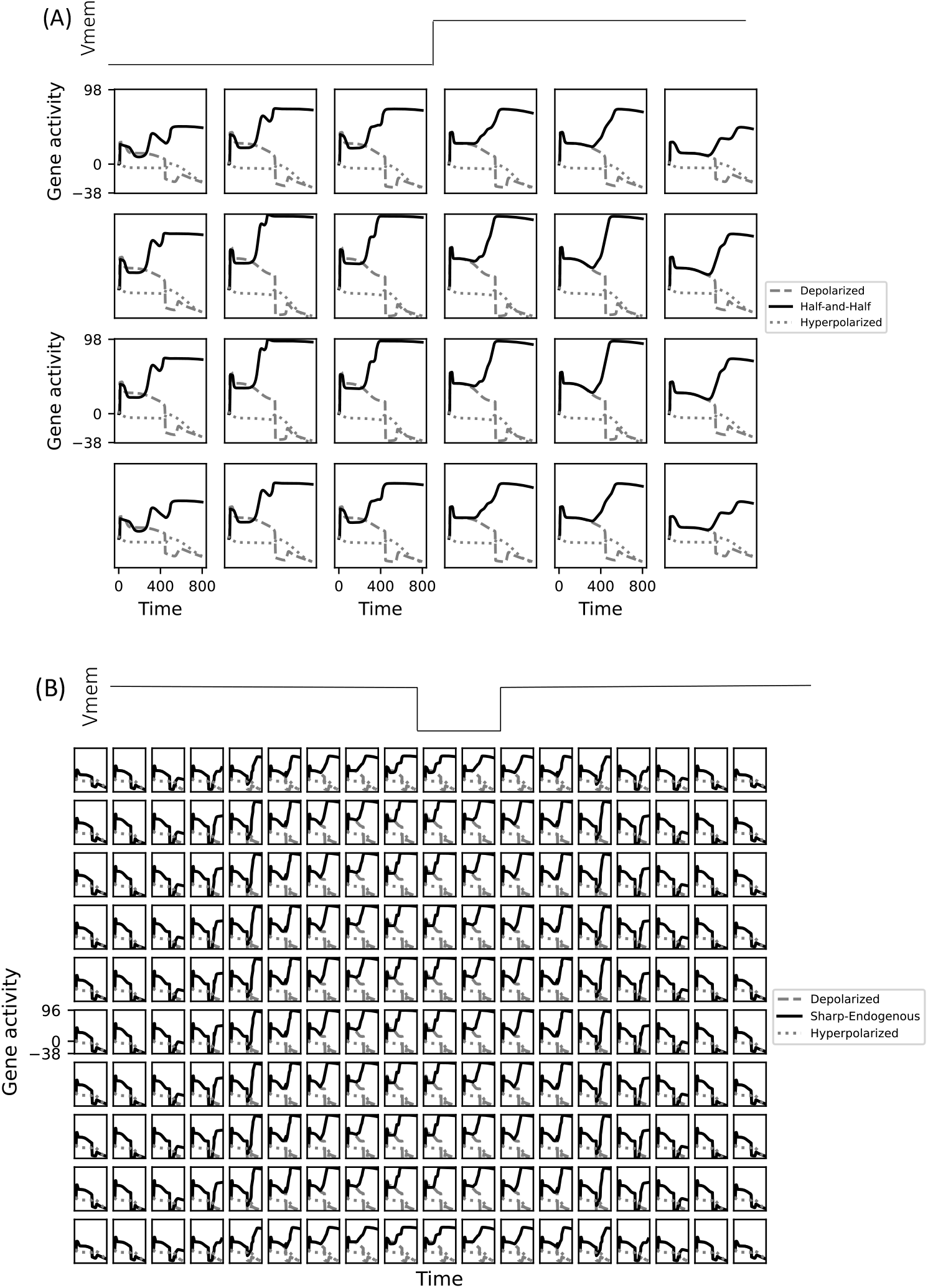
Cell-wise behavior of the discriminator gene while responding to aberrant bioelectric input patterns: (A) A half-and-half Vmem pattern where the left half is hyperpolarized, and the right half is depolarized (inset) results in normal brain development as indicated by the activation of the discriminator gene in all cells, compared to deactivation for the uniform patterns. (B) A sharpened endogenous Vmem pattern where only a third of the hyperpolarized central band of the endogenous pattern is hyperpolarized (bottom; the middle two columns of the six at the center of this 10×18 tissue) results in an intermediate brain development due to the failure of gene expression to discriminate between the endogenous and the incorrect Vmem patterns in the leftmost and the rightmost two columns.

### 9. The inclinations of context-dependency of the cells in a 4×6 tissue while solving the pattern discrimination problem

**Figure S6.**
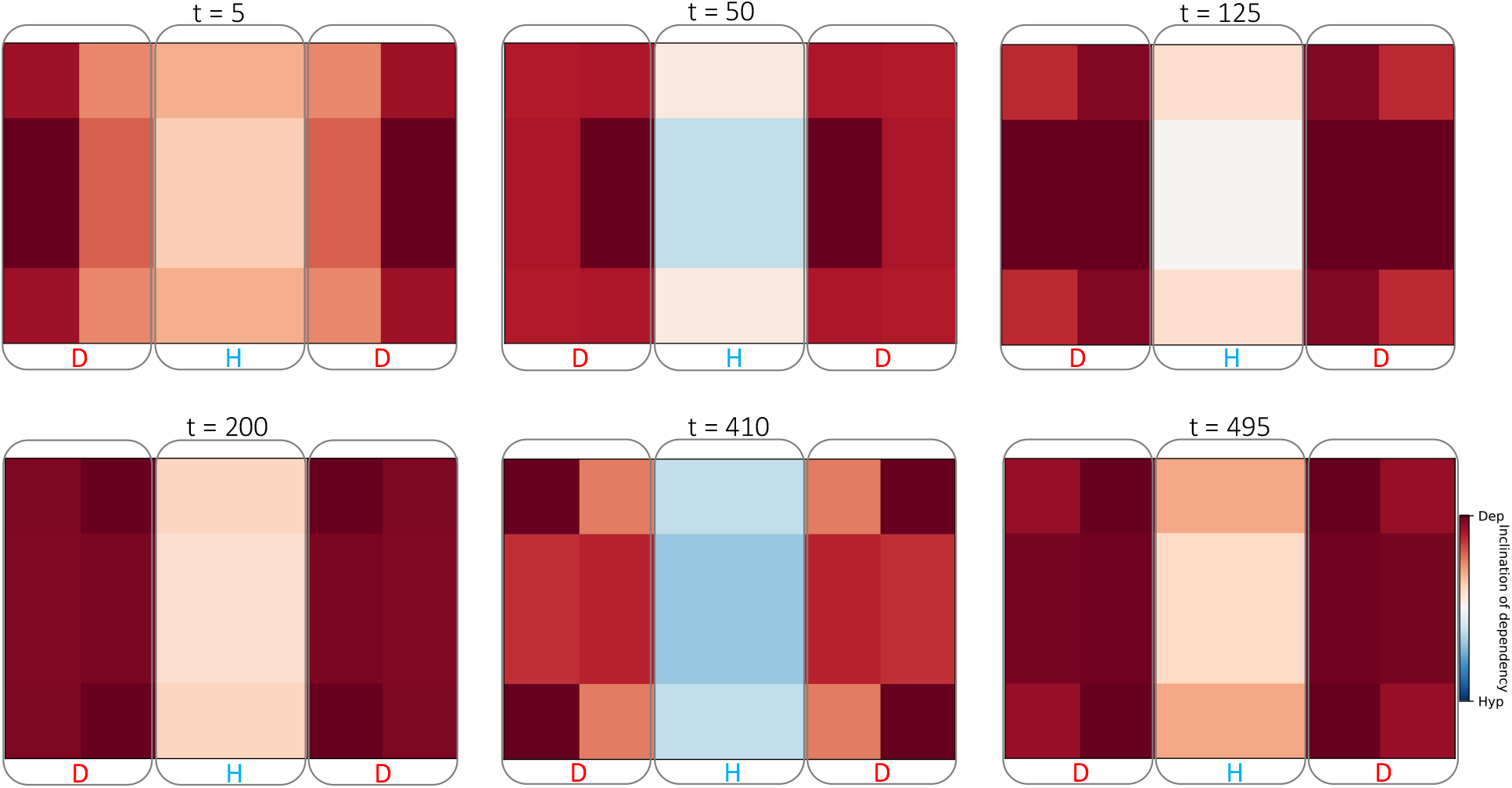
The time-evolution of context-dependency of the individual cells in a 4×6 tissue while solving the pattern discrimination problem. The color of each cell indicates the extent to which the connections of that cell in the corresponding Hessian network originate from another depolarized cell in the left or the right bands or a hyperpolarized cell in the central band.

### 10. The behavior of a canalized simple recurrent dynamical system can be reconstructed using partly linearized Taylor expansions involving the control parameter

To help the reader better understand the concept of canalizing dynamics in our complex neural plate model we illustrate it in an analogous simple recurrent ODE model. The main idea is that a nonlinear recurrent system could effectively act as a relatively linear feedforward-like system if the coupling between the controlling parameter (external input) and the controlled variable is set up in such a way that the former dominates the latter.

Consider the following ODE consisting of a single recurrent variable *x* controlled by parameter *a*:

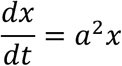

Even though this equation can be analytically solved as 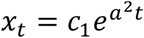, where *c*_1_ is the integration constant, we shall consider the approximated timeseries *x*_1_, *x*_2_, …, *x_t_*, … generated by Euler method:

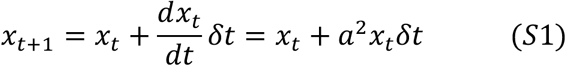

The above can also be expressed in equivalent “parameter-Taylor” (PT) expansions consisting of the terms 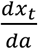, representing the sensitivity of the variable to the controlling parameter. This shall allow us to understand the role of the parameter in reconstructing the timeseries of the variable *x*. For example,

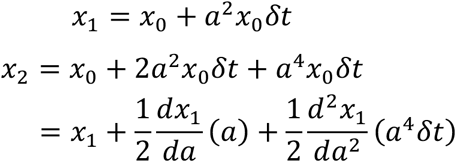

Where:

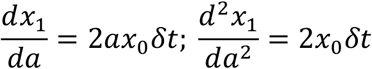

Likewise,

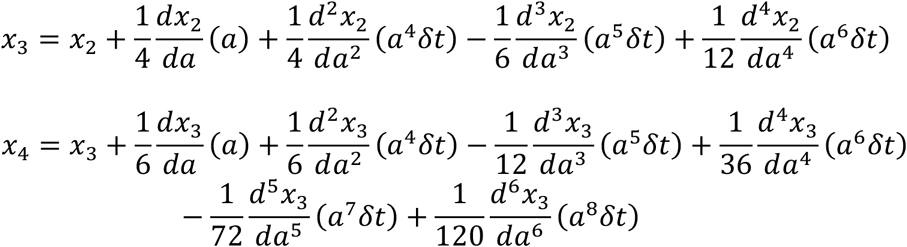

And so on.

Given that Eqn. S1 can be rewritten as *x_t_* = *x*_0_(1 + *a*^2^*δt*)^*t*^, the derivatives of *x_t_* with respect to *a* can be expressed in generic terms as follows:

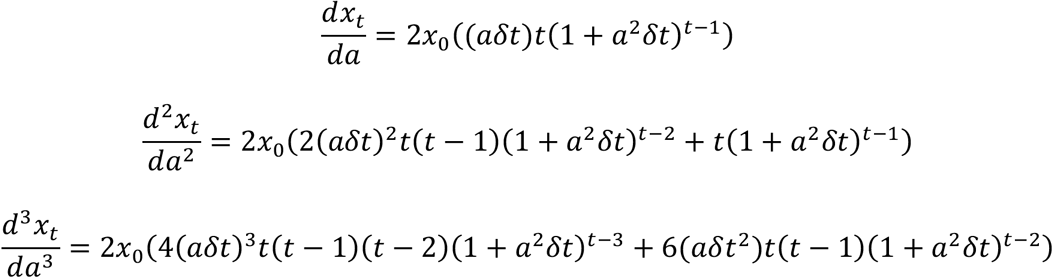

And so on.

As can be noticed, the coefficient of the leading term is given by, 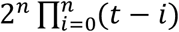, where *n* refers to the order of differentiation. Clearly, this coefficient grows nonlinearly with *t* and *n*. This is the reason why the coefficients of the derivatives in the PT expansions shrink nonlinearly (to compensate for the nonlinear growth of the derivatives) with time for the higher order derivatives, as can be seen in the expressions above. How informative are these coefficients in reconstructing the timeseries of the variable? Is their nonlinear character indispensable for that purpose?

Even though it may seem that time-dependent PT coefficients are required to reconstruct the timeseries of the variable, below we show that static coefficients are sufficient for a qualitative reconstruction. We also show that good reconstructions can be achieved even with just the 1^st^ and 2^nd^ order Taylor terms. We’ll see that this is possible because of the monotonic, even if nonlinear, character of *x*(*t*).

We reconstruct the observed timeseries 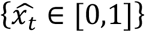 obtained by normalizing the raw timeseries {*x_t_, x*_0_ = 1}, by fitting a “pruned timeless parameter-Taylor” (PTPT) equation, where the 1^st^ and 2^nd^ order derivatives are known, and the time-independent coefficients *c*_1_ and *c*_2_ are optimized using gradient descent:

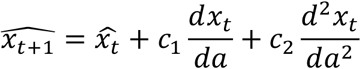

Note that we ignored the terms (*a*) and (*a*^4^*δt*) corresponding to the 1^st^ and 2^nd^ order derivatives since they are constants. Here, we can observe that the reconstruction is better for a larger parameter value (*a* = 10) compared to that obtained for a smaller parameter (*a* = 1). This follows a general trend, where the reconstruction error tends to get smaller for larger parameter values (Fig. S2). These observations suggest that “timeless” Taylor coefficients are sufficient to reconstruct an effectively feed-forward-like system (large α), that is a system where the external input effectively canalizes the dynamics of the variable. This is because a canalized system reduces the nonlinearity of the system by partly linearizing it – an effect of the leveraging of timeless Taylor coefficients.

We leverage this observation to hypothesize that a dynamical model, such as our neural plate model, whose PTPT reconstructions yield good fits effectively acts like a simplified canalizing system such as the example described here.

**Figure S7.**
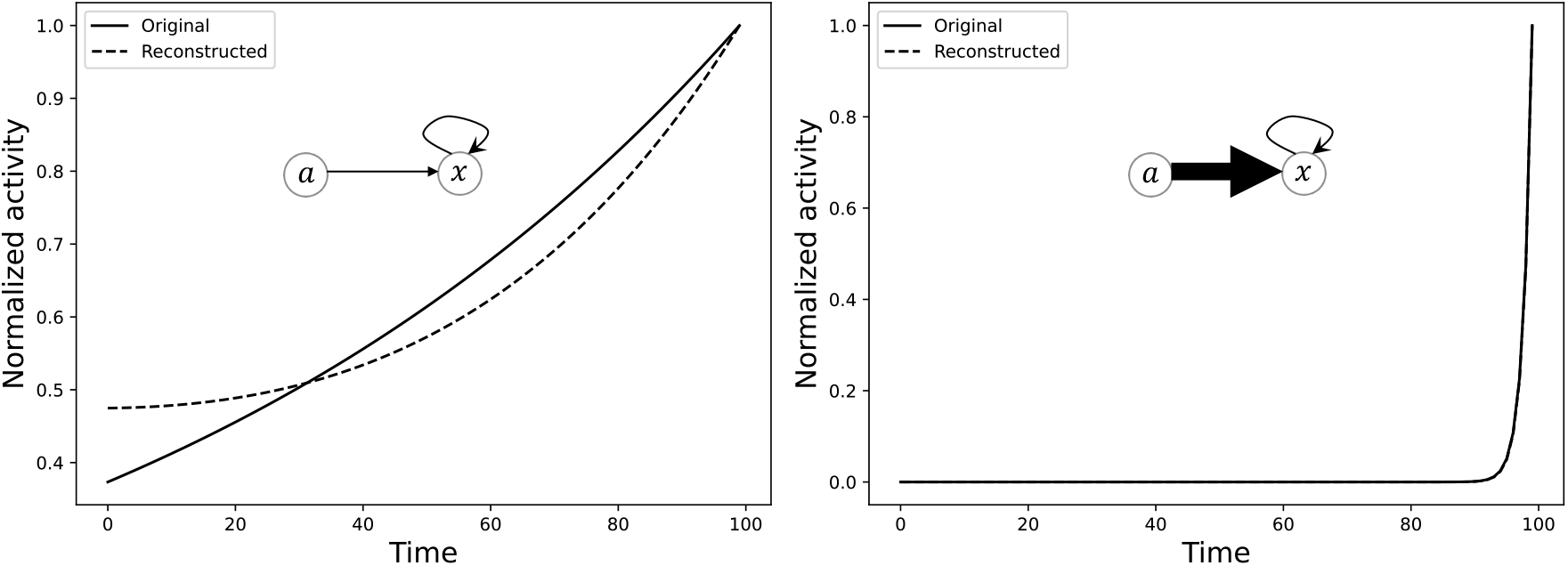
A nonlinear recurrent dynamical system can effectively act like a linear feed-forward-like system when the forcing parameter controls the variable more strongly than the variable itself. Shown here are reconstructions of the original observed timeseries using a PTPT expression, where (left) the parameter is small (*a* = 1); and (right) the parameter is relatively large (*a* = 10).

**Figure S8.**
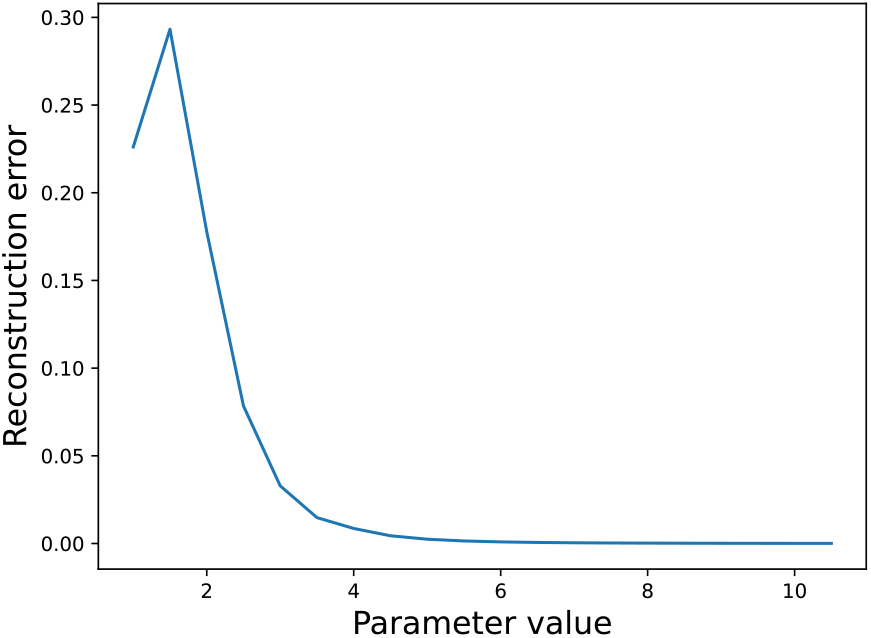
The quality of the simplified Taylor reconstructions with time-independent coefficients (PTPT) gets better with more canalization (larger values of the forcing parameter *a*). The reconstruction errors were computed as the mean squared error between the reconstructed and the original timeseries.

### 11. A strong higher-order relationship between the spatial resting potential and gene expression patterns cannot be expected in random models

We fit a PTPT equation to reconstruct the normalized gene expression timeseries for a suite of 100 randomly connected parametrized neural plate models by optimizing either the Jacobian coefficient *c_J_* or the Hessian coefficient *c_H_*, both vectors of length 1 × *n_g_*.

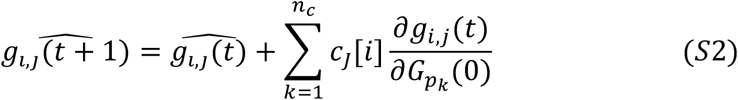

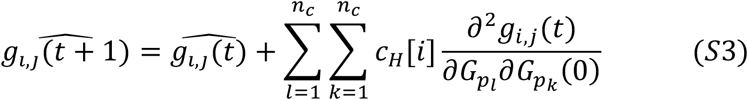

Here, 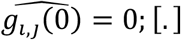 refers to the gene index. The distribution of the Jacobian reconstruction errors (MSE) thus obtained by optimizing Eqn. S2 suggests that good fits should be expected (Fig. S3), with the 5-percentile MSE being as low as 0.07. Moreover, the Jacobian reconstruction MSE of our neural plate model is about 0.14 and is well placed within the expected range of 5 and 95 percentiles. The Hessian reconstruction errors obtained by optimizing Eqn. S3, on the other hand, are higher (Fig. S3) than the corresponding Jacobian errors (5-percentile MSE is about 4.8). Although, the Hessian error of our neural plate, with an MSE of 0.23) is well below the expected range.

These observations suggest that the neural plate model was optimized to operate at the second-order level.

**Figure S9.**
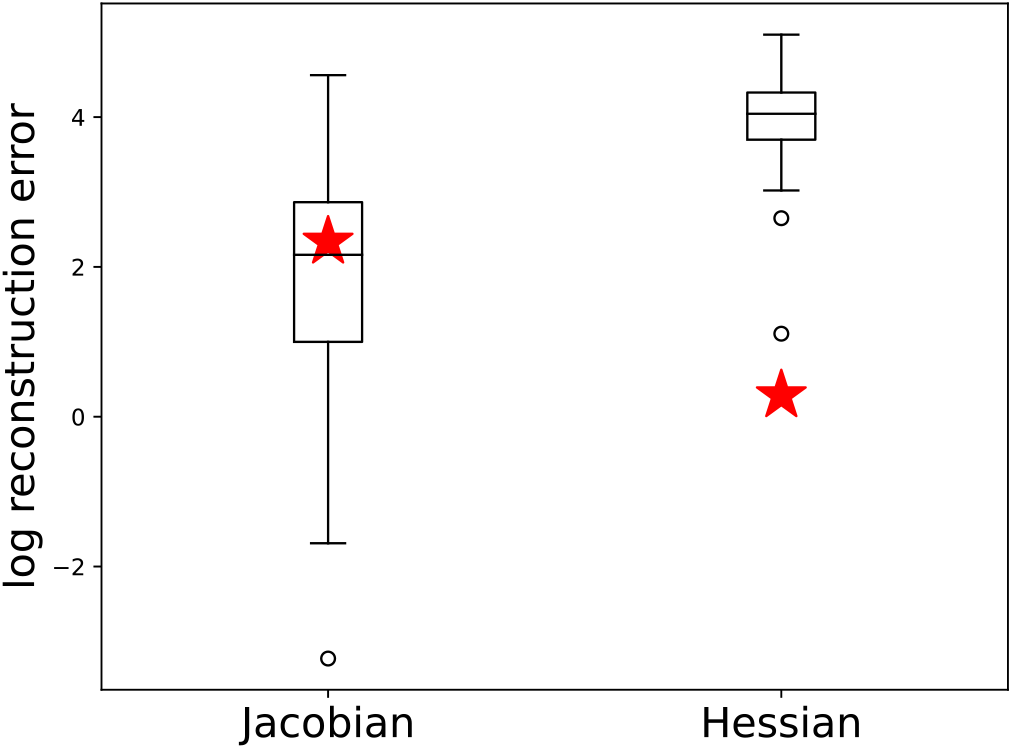
The distributions of reconstruction errors, plotted on a log scale, for a suite of 100 randomly parametrized neural plate models obtained by individually optimizing the coefficients of the Jacobian and the Hessian terms. Red stars indicate the corresponding errors for the trained neural plate model described and analyzed in the main text.

### 12. Mathematical expressions behind the quantities used in Figs 8–10

Fig. 8 depicts sum of the sensitivities of the discriminator gene (*g*_6_), corresponding to the endogenous input pattern, in the cells in just the top left quadrant of the tissue, which is given by:

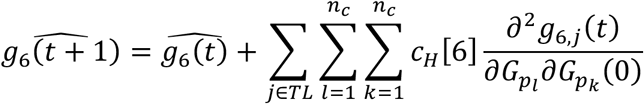

where:

*c_H_* (a vector of length 1 × *n_g_*) is optimized to maximally fit the reconstructed 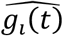 with the observed *g_i_*(*t*);
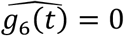; and *TL* refers to the collection of cells in the top left quadrant of the tissue.

Fig. 9 is a depiction of the following quantity, where *t* represents the time points indicated in the figure, and the other variables are as described in the equation above:

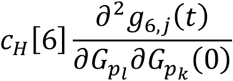

Fig. 10 depicts the net scale sensitivity, *S(i*), of each gene, where:

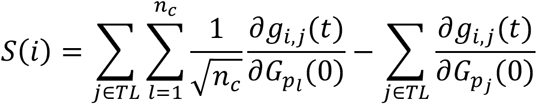

Here, the first term is a directional derivative and computes the net sensitivity to the tissue, and the second term refers to the single cell. The term 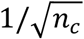 is a normalization term for computing the directional derivative along the direction of the unit vector, thus enabling the comparison between the two scales of the tissue and the single cell.

### 13. A higher-order and canalization-like mechanism characterizes the relationship between the spatial Vmem pattern and gene expression

**Figure S10.**
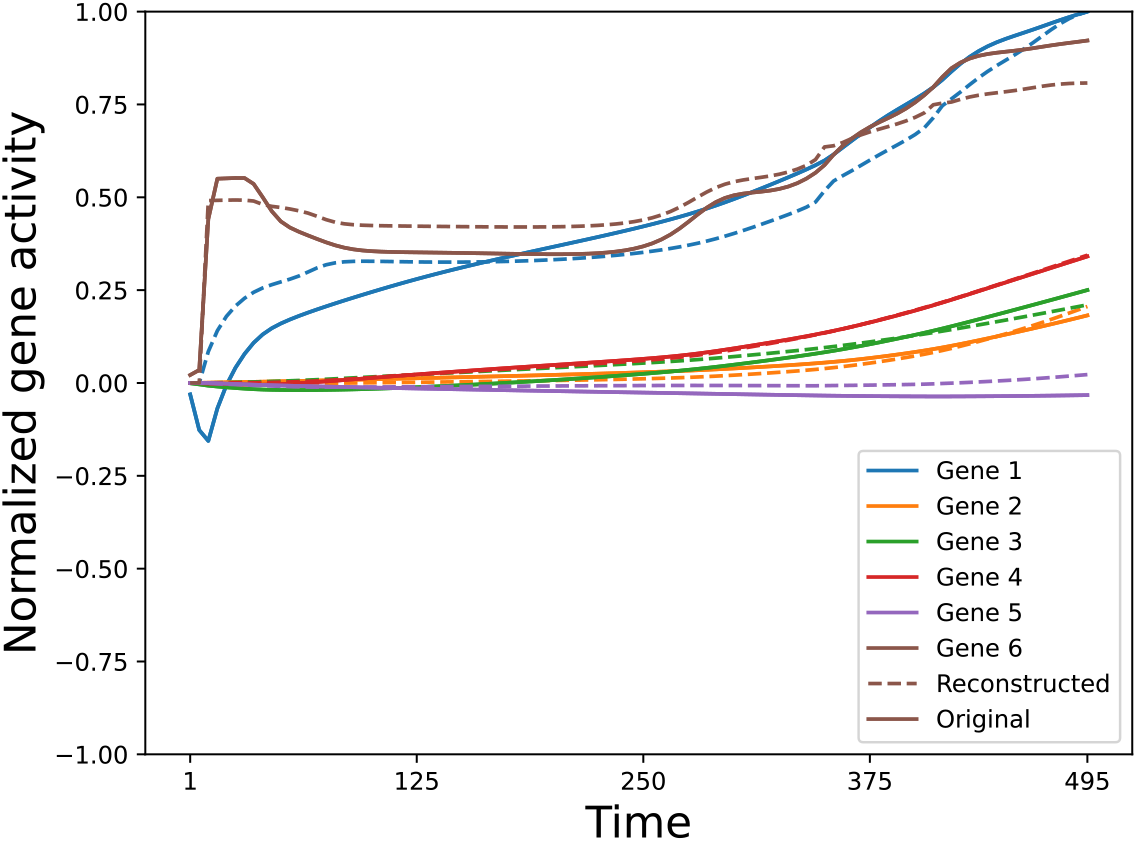
Gene-wise Hessian reconstruction of timeseries. A comparison of the normalized original gene expression timeseries of all the genes (dashed) and the corresponding reconstruction (solid) from the second order (Hessian) influence of the Vmem. The resemblance between the reconstruction and the original indicates that most of the gene activity is determined by its second-order relationship with Vmem. This moreover also suggests a canalization-like mechanism where the dynamics of this complex recurrent model is simplified in such a way that there is an almost direct control of the discriminator gene by the Vmem, as if the paths through the other genes are bypassed; this canalized pathway cannot be inferred from the structure of the static connectivity (Fig 3) alone. This makes intuitive sense since discriminatory behavior requires a comparison of Vmem of different cells, which the Hessian presumably aides by containing the differential information. Due to the symmetry of the model, only the genes contained by the cells in the 2×3 top left quadrant of the full 4×6 tissue (dashed boxes in Fig 9) were considered for these calculations; the Hessian depicted here is the sum of all Hessians corresponding to each cell in the quadrant.

### 14. An oscillatory scanning-like behavior characterizes bioelectric control

The discriminator gene (Gene 6) exhibits a unique pattern in its Jacobian where it displays an oscillatory scanning behavior by constantly shuffling the order of its sensitivities to the three resting potential pattern bands (Fig S10). In particular, the gene shuffles its sensitivity to the three bands through a sequence of five distinct orderings (Figure S10A) in comparison to three orderings by the average gene (Figure S10B). Such a shuffling strategy may be employed as a means to integrate information from the three resting potential pattern bands before making a final decision.

**Figure S11.**
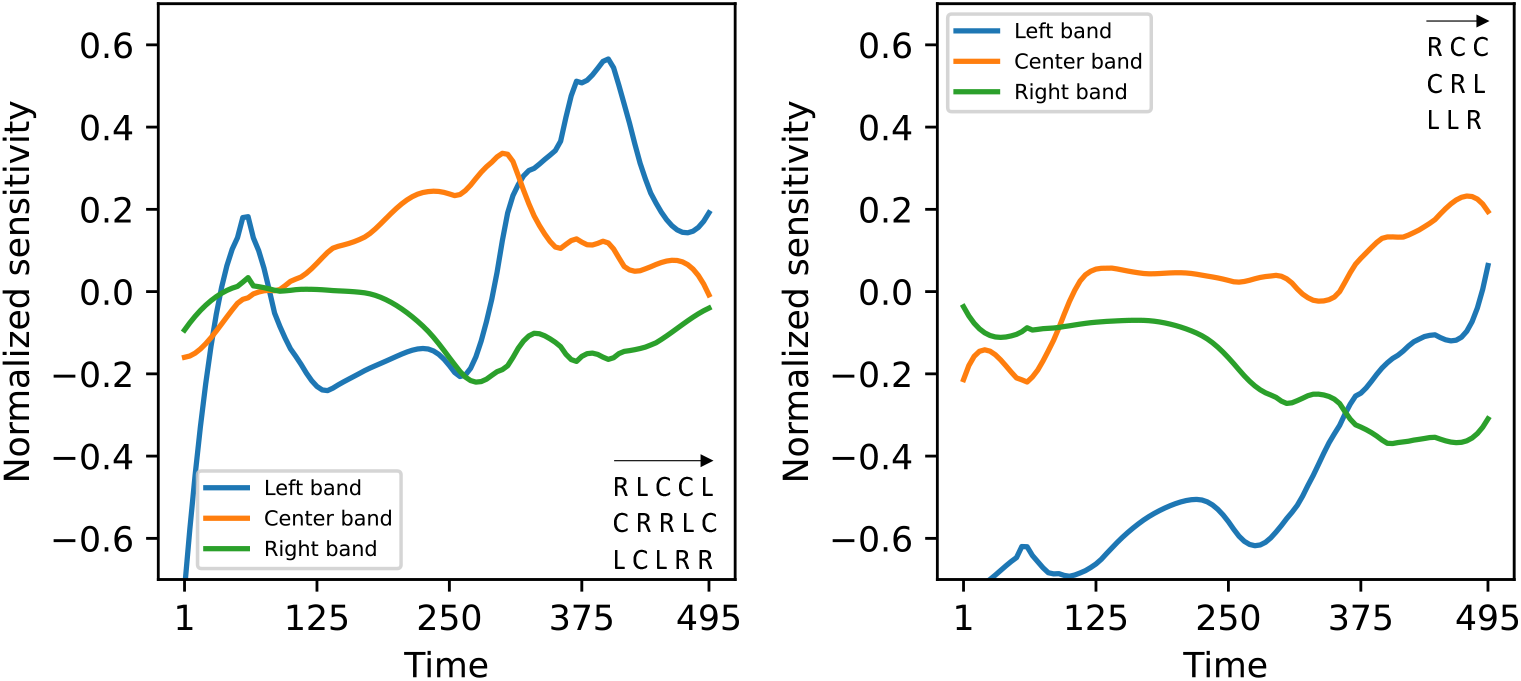
The discriminator gene dynamically processes the three resting potential pattern bands (Figure 1F) in a more complex way (**A**) compared to the average gene (**B**). The band-wise first order (Jacobian) normalized sensitivities of gene activity computed with respect to the resting potential pattern bands has a uniquely complex profile for the discriminator gene-gene 6 (**A**) where the ordering of the bands in terms of their signed sensitivities goes through a slightly more complex sequence (bottom right inset) compared to the average gene (**B**). From the perspective of minimal cognition, one could conceive this as a form of shifting of “attention”, where attention is constantly shifted from one band to the other so that the corresponding information could be captured and then integrated before settling on a decision. Due to the symmetry of the model, only the cells in the 2×3 top left quadrant of the full 4×6 tissue were considered for these calculations.

### 15. Different genes play different roles in solving the patterning problem

**Figure S12.**
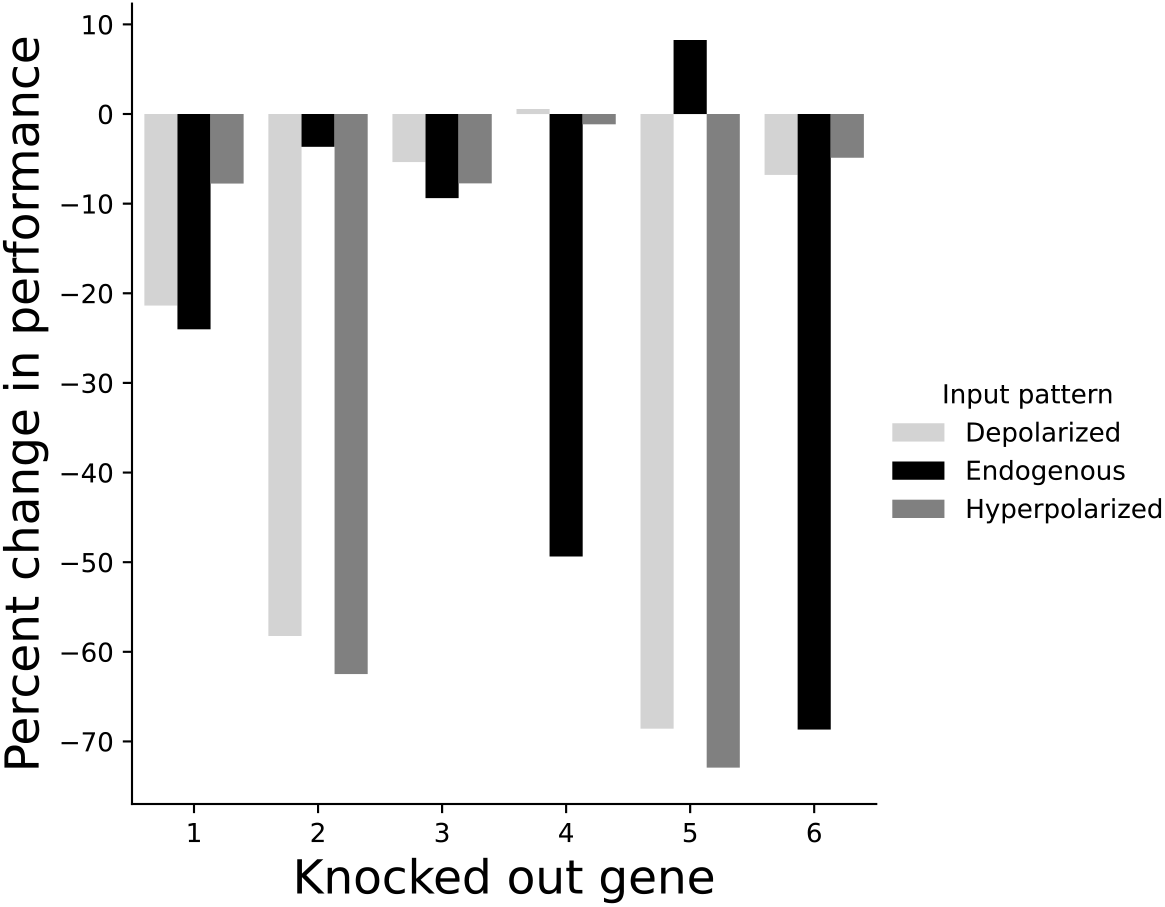
Differential effect of gene knockouts on the performance of the model for each of the three input bioelectric patterns. It can be noticed that gene 6 effects the largest drop in performance for the endogenous pattern while contributing very little to processing the other patterns. While gene 4 behaves in a similar fashion as gene 6, genes 2 and 5 behave conversely where they most affect the depolarized and the hyperpolarized pattens while contributing very little to the endogenous pattern. Gene 3 appears to be the most redundant gene of the lot by affecting very little the performance with respect to all three patterns.

### 16. More examples of scale robustness

**Figure S13.**
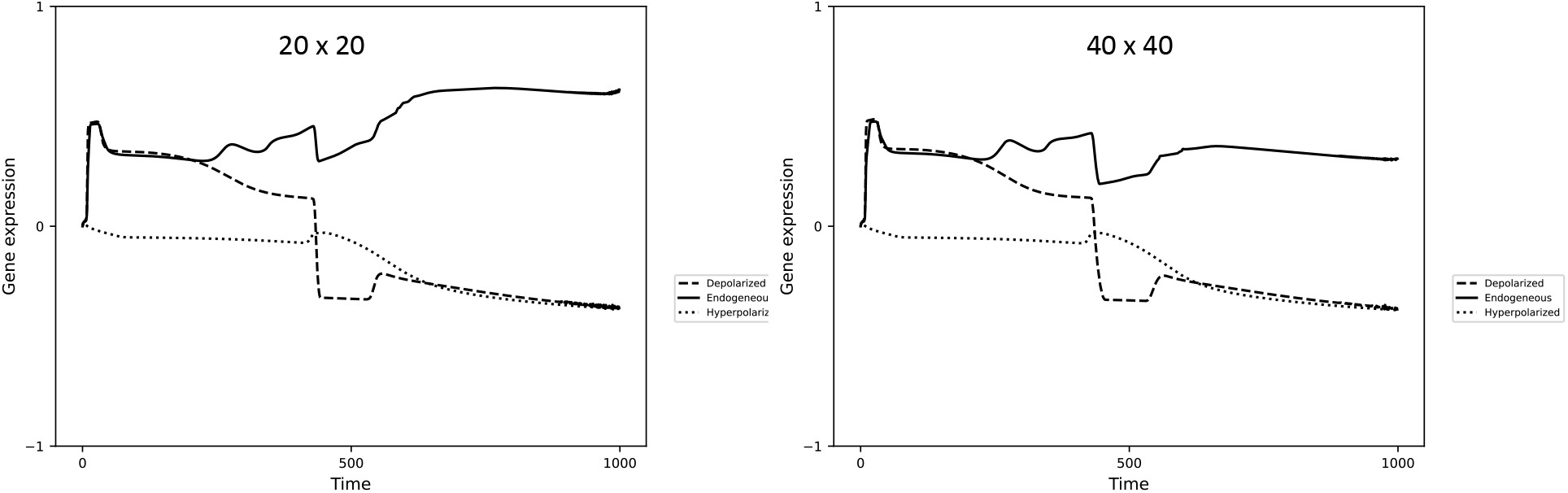
The model solves the pattern discrimination problem for tissues of size 20×20 (left) and 40×40 (right). Shown here is the timeseries of the discriminator gene. The corresponding gene expression timeseries are qualitatively similar to that of the original tissue of size 4×6 for which it was trained (Fig 5A inset).

### 17. Severing the connections from the genes to the Vmem impedes performance

**Figure S14.**
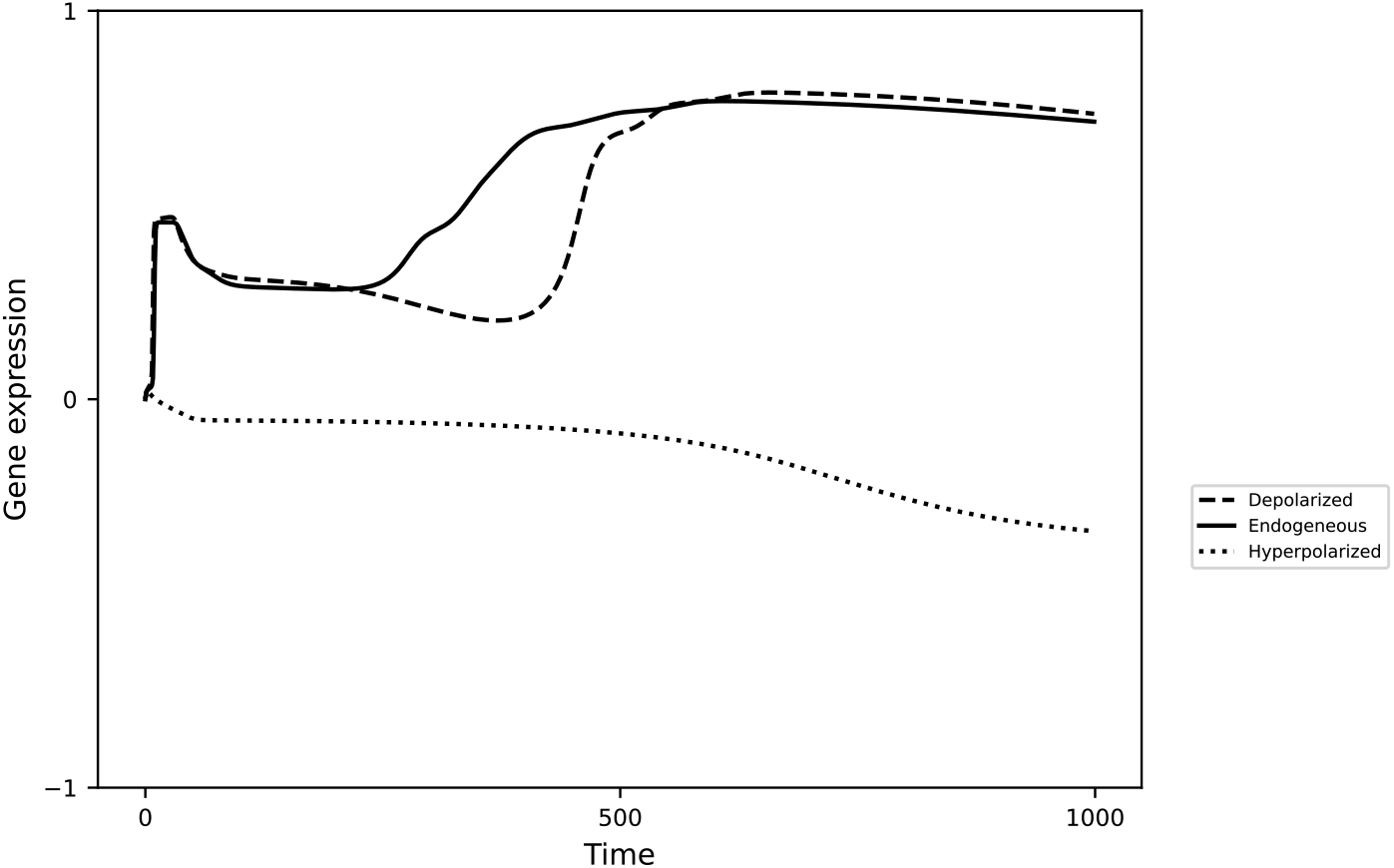
The timeseries of the discriminator gene following the severing of the connections running from the genes to the Vmem in all cells. It can be seen that the detection of the depolarized voltage pattern is affected, though the behavior with respect to the other patterns are preserved.

